# Cell-autonomous differentiation of human primed embryonic stem cells into trophoblastic syncytia through the nascent amnion-like cell state

**DOI:** 10.1101/2021.06.28.450118

**Authors:** Masatoshi Ohgushi, Mototsugu Eiraku

## Abstract

Human primed embryonic stem cells (pESCs) are known to be converted to cells with several trophoblast properties, but it has remained controversial whether this phenomenon represents the inherent differentiation competence of human pESCs to trophoblast lineages. In this study, we report that chemical blockage of ACTIVIN/NODAL and FGF signals is sufficient to steer human pESCs into GATA3-expressing cells that give rise to hormone-producing syncytia analogous to syncytiotrophoblasts of the post-implantation stage of the human embryo. Taking advantage of this system, we identified two distinct modes of cell-autonomous genetic programs and their coordinated actions to initiate the differentiation. We also found a transient population reminiscent of nascent amnion and then a spontaneous branch of differentiation trajectory leading to syncytiotrophoblast-like syncytial cells. These results provide insights into the possible extraembryonic differentiation pathway that is unique in primate embryogenesis and is relevant to the trophoblast competence of human primed pluripotent stem cells.

## Introduction

In mammalian development, a totipotent fertilized egg undergoes a series of cell divisions while differentiating into several lineages in a stepwise manner. The intrinsic genetic program and cell-cell communication coordinate to control these processes by instructing progenitor cells on each developmental path. Genetic studies using mice as model animals have established the concept that the first cell fate segregation in mammals, the inner cell mass (ICM)- or trophectoderm (TE)- lineage segregation, occurs around the morula stage. These fate-committed cells produce a well-organized blastocyst at a subsequent developmental stage. ICM is a transient pluripotent tissue that gives rise to all cells of the fetus, while TE provides multiple components of the placenta that plays important roles in implantation and fetal growth. Several lineage-tracing studies revealed that the developmental paths of ICM- and TE-progenies never crossed once their identities were established (reviewed in Rossant, 2018). Consistent with the developmental dogma, ICM-derived ESCs retain pluripotency, and therefore, can differentiate into cells of all three germ layers (Martello and Smith, 2014). When injected into blastocysts, ESCs are incorporated into ICM tissue and start to differentiate following the developmental program of the host animals. In chimeric pups, ESC derivatives are distributed in multiple tissues, except for the placenta. This provides strong evidence that mouse ESCs lose the competence to differentiate into trophoblast lineages. Indeed, trophoblast differentiation from mouse ESCs is not attainable *in vitro* unless artificial genetic manipulations are conducted.

The consensus that ESCs never differentiate into trophoblast lineages has been challenged by a surprising report showing that human ESCs were converted into differentiated cells with unique properties for placental cells just by exposing them to BMP4 (Xu et al., 2002). Many groups succeeded in replicating this phenomenon; in particular, the establishment of a protocol to achieve the efficient conversion of human pESCs into trophoblasts, called a BAP (BMP4 plus activin blocker A83-01 and FGF blocker PD03714) facilitated the reproducibility (Amita et al., 2013; Roberts et al., 2018). The pESC-derived trophoblast-like cells has been utilized as an experimental platform for molecular and pathological studies of viral infection or placental disorders, and provide valuable information (Sheridan et al., 2017; Sheridan et al., 2019; Horii et al., 2021). Nevertheless, the reliability of the pESC-to-trophoblast differentiation has been under intense debate over the past decades (Bernardo et al., 2011; Roberts et al., 2014), possibly because it contains a striking contradiction to the developmental dogma that has been validated in mouse studies. Recently, two groups claimed that the BMPs induce amnion rather than trophoblast differentiation in human pESCs, questioning their trophoblast competence (Guo et al., 2021; Io et al., 2021). Whether trophoblast-like differentiation from pESCs recapitulates any developmental processes remains an open question.

During the investigation of BAP-induced responses, we noticed that without exogenous supplementation of BMP ligands, human pESCs differentiate into cells expressing several extraembryonic marker genes, including GATA3. This finding suggests that when an undifferentiated state is destabilized, pESCs direct into GATA3- expressing extraembryonic lineages in a cell-autonomous manner. Using this differentiation system, we sought to characterize the cellular properties of pESC-derived GATA3^+^ cells and deciphered the molecular mechanisms underlying the cell state transition linking pluripotency dissolution to the GATA3^+^ cells. Our results show that human primed pluripotent stem cells have the competence to differentiate into hormone-producing syncytial cells through an amnion-like cell state.

## Results

### ACTIVIN/FGF blockers alone are sufficient to generate GATA3^+^ cells from human pESCs

To investigate the dynamic process of trophoblast-like differentiation from human pESCs, we generated reporter cell lines by applying GATA3 as a readout for the trophoblast identity (G3KI cells, clones #5-2 and #30-1, Figures S1A-B). We confirmed that these reporter lines became positive for tandem tomato (tdT) upon a BAP treatment (Figure S1C).

Interestingly, during the validation of these reporter lines, we noticed that exogenous BMP4 was not essential for producing GATA3-expressing cells (Figure 1A). Flow cytometric analyses showed that > 95% of cells expressed tdT at day 3 just by adding only two inhibitors (Figures 1B-C, S1D). On day 3, tdT fluorescence showed two distinct peaks, but became uniform by day 6. These tdT^+^ cells also expressed a panel of trophoblast-associated markers (Figures 1D-F). Quantitative RT-PCR analyses confirmed that genes associated with placental functions, including hormone biosynthesis, nutrient transport and immunological tolerance, showed a significant increase in their mRNA expression (Figures 1G, S1E). As different placental features, the upregulation of endogenous retroviruses (ERVs), the syncytialization, and the expression and/or secretion of placental hormones chorionic gonadotrophin β (CGβ), GDF15 and PGF, were detected after ECSs were treated with the two inhibitors (Figures 1G-J). To monitor syncytium formation, two lines of AP-treated pESCs, each of which was marked by different fluorescence proteins (H2B-Venus and mCherry, respectively), were mixed and then co-cultured. After 4-6 days, we detected the emergence of many syncytia, which were defined by the co-expression of both fluorescent proteins (Figures 1K-L, S1F).

**Figure 1.**
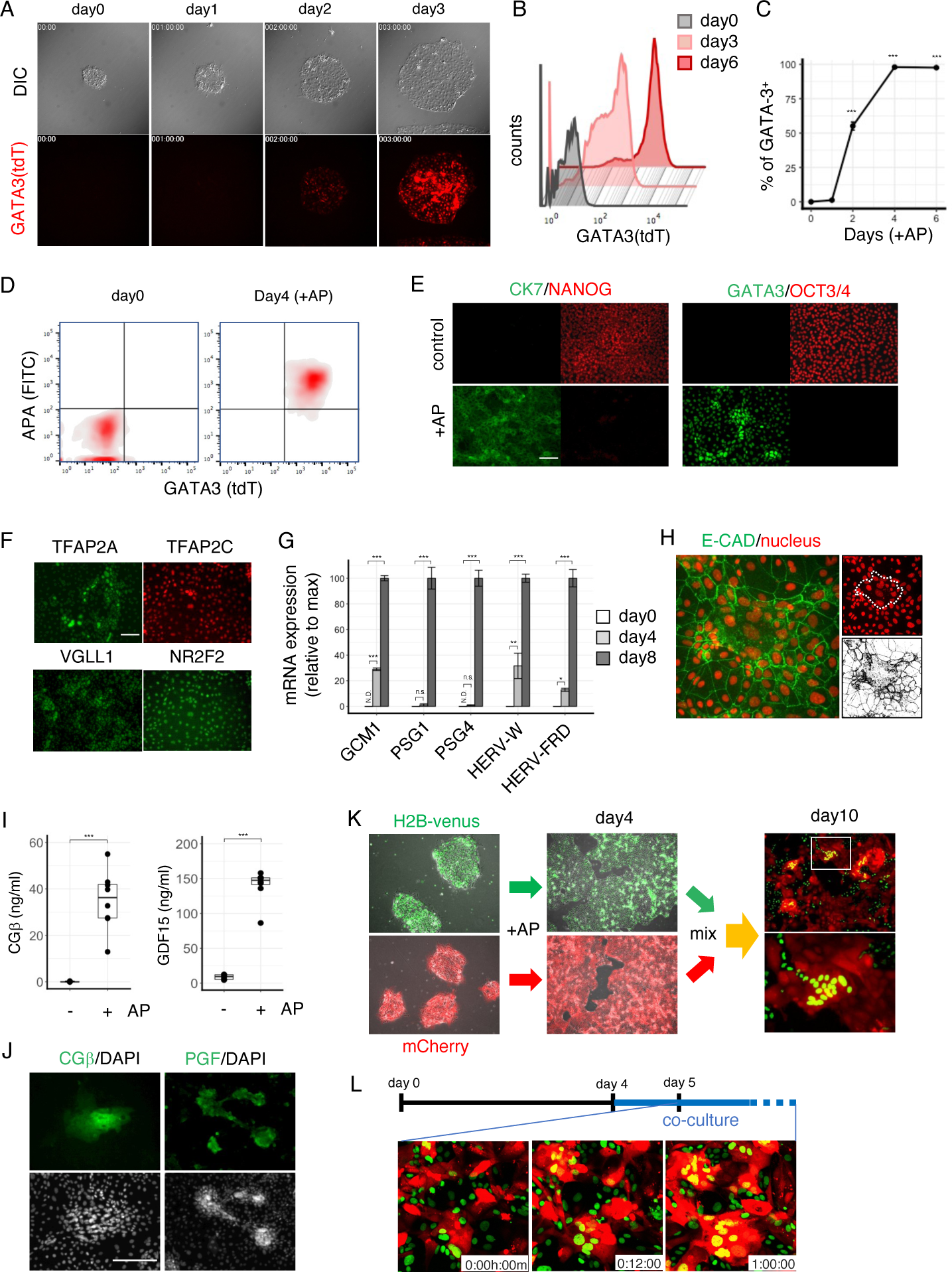
ACTIVIN/FGF blockers alone are sufficient to generate GATA3^+^ cells from human pESCs. (A) Live imaging of the GATA3-knockin cells treated with AP for 72 h. Red fluorescence indicates GATA3-expressing cells. Recording was stared immediately after AP addition. (B, C) Time course quantitation of GATA3^+^ cells after AP addition. (D) Flow cytometric panels for APA^+^/GATA3^+^ cells at day 4 after AP addition. (E) Immunostaining for pESC or trophoblast markers before and 3 days after of AP addition. (F) Immunostaining for trophoblast markers 4 days after of AP addition. (G) qPCR assay for trophoblast-related genes. GATA3-knockin cells were treated with AP for the indicated periods. (H) Confocal images of syncytia at 4 days after AP addition. A binary image of ECAD- signal (right, gray image). Syncytia is defined with E-CAD signal, and indicated as the enclosed regions in the right fluorescence image (dotted lines). (I-J) Production of placental hormones. Secretion level of CGβ and GDF15 in AP-treated cells (at day 8) (I). Immunostaining for CGβ and PGF at 6 days after AP addition (J). (K-L) Visualization of syncytium formation. Syncytia were double-positive for both fluorescence proteins (K). Time lapse conformal images of syncytialization (L). Coculture was started at 4 days after differentiation, and fluorescent images were recorded from day 5. Scale bars show 200 µm. Error bars in the graphs represent standard deviations. Dunnett’ test (n = 3) versus control (day0) (C, G); Wilcoxon signed-rank test for paired data (H); N. D., not detected, n. s., not significant; \**p* < 0.05, ** *p* < 0.01, *** *p* < 0.001.

Taken together, these observations indicate that the combined inhibition of NODAL/ACTIVIN and FGF signals steered pESCs to express biological features unique to the placenta. We observed GATA3 induction in all culture media and substrates we tested and in other primed ESC or iPSC lines (Figures S1G-J). Hereafter, we denote this differentiation recipe consisting of small molecules only as an ‘AP’ (BAP minus BMP4), and GATA3-expressing cells as ‘GATA3^+^ ExE cells’ to distinguish them from trophectoderm-derived conventional trophoblast lineages.

### Transcriptome characterization of AP-induced GATA3^+^ cells

We explored the genome-wide transcriptome features and dynamics of AP-induced differentiation processes. Principal component and clustering analyses indicated a drastic change in gene expression patterns between 2 and 4 days after AP addition (Figures 2A, S2A). These differentiated cells hardly expressed markers for definitive endoderm, mesoderm and neural progenitors (Figure S2B, Cliff et al., 2017). We then extracted differentially expressed genes (DEGs) between pESCs and GATA3^+^ ExE cells (at day8) and performed gene enrichment analyses (Figure 2B). These analyses indicated that the upregulated genes were more closely associated with the placenta than other tissues and with placenta-related terms or biological processes (Figures 2C, S2C). These results indicated that AP evoked differentiation into cells that were transcriptionally assigned to the trophoblast or placenta.

**Figure 2.**
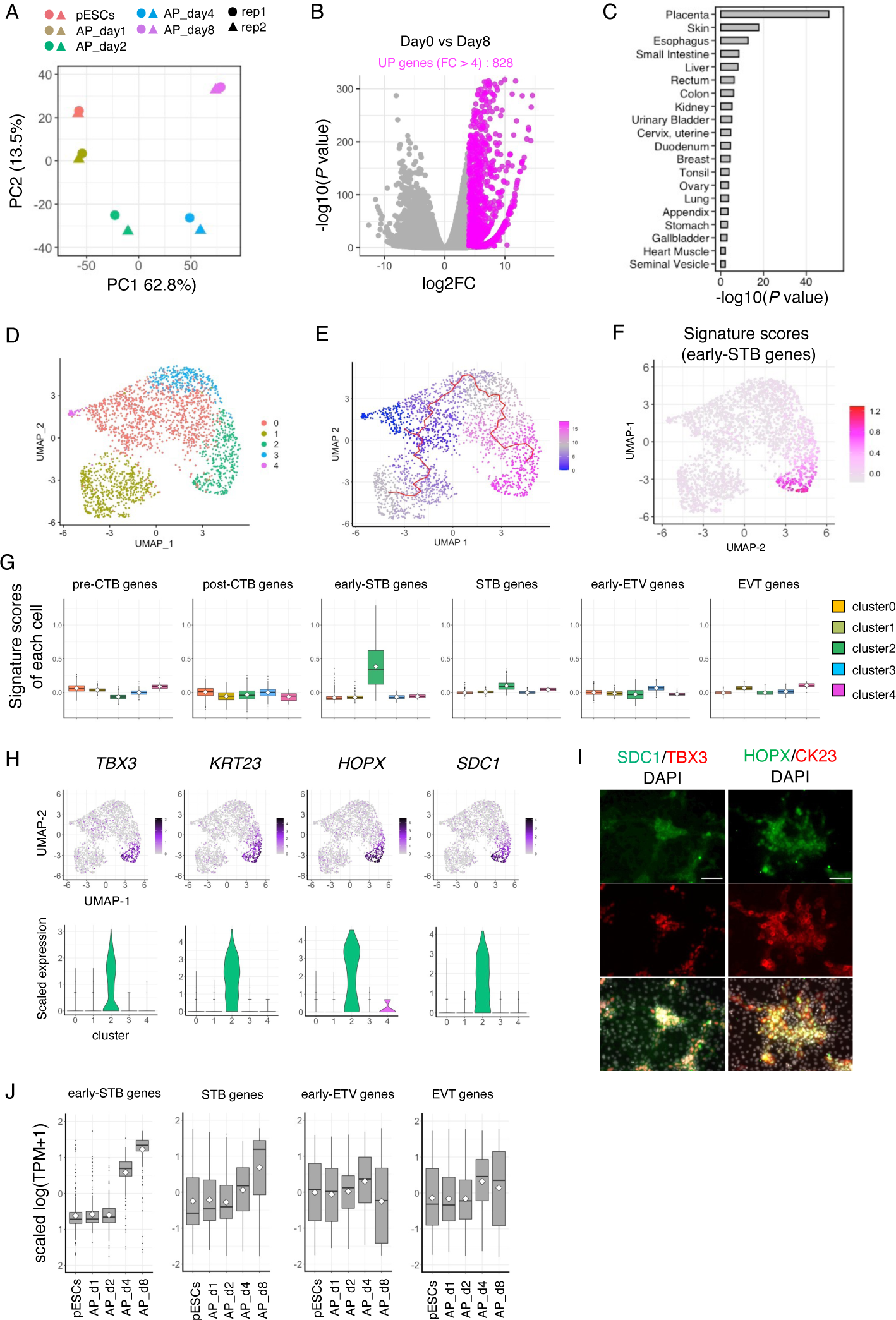
Transcriptome characterization of AP-induced GATA3^+^ cells. (A) PCA analyses of AP-induced differentiated cells at day 0, 1, 2, 4 and 8. The top 1,000 valuable genes were used for the variation calculation. (B) Volcano plot of pESCs *vs* differentiated cells (day8). The upregulated genes (n = 828, log2FC > 4, *p*-value > 0.05) are indicated as red dots. (C) Tissue distribution of the identified genes (n = 721). (D) UMAP representation of 2,464 AP-induced GATA3^+^ cells (day 4). The five clusters can be visualized based on the colors indicated. (E) Cellular trajectory reconstruction using Monocle3 applied onto cell clusters in a UMAP space. Color gradient represents the transition of estimated pseudotime, and red lines show the estimated trajectory. (F) Signature score distribution at day 4 of differentiation. Signature scores were calculated using genesets for the early STB genes (n = 218) and applied onto cell clusters in a UMAP. (G) Boxplot representation of the signature scores for each cell. Signature scores were calculated using geneset for each trophoblast lineage that was defined by Zhang et al. (2019) (For pre-CTB, n = 213; post-CTB, n = 97; E-STB, n= 218; STB, n = 222; E-EVT, n = 148; EVT, n = 395). Data are shown grouping with the indicated clusters. (H) Scatter and violon plot representations of the cluster 2-enriched genes identified using scRNA-seq data. (I) Immunostaining using KhES-1 cells for the cluster 2-enriched genes at day 5 after AP addition. (J) Boxplot presentation showing the expression dynamics of the indicated trophoblast marker genes based on bulk-mRNA-sequencing data. The scaled TPM values for E-STB (n= 207), STB (n = 2,204), E-EVT (n = 150) and EVT genes (n = 365) are shown. Scale bars show 100 µm.

Recently, gene expression signatures of human peri-implantation embryos from 6 to 14 days post-fertilization (d.p.f) were described at the single-cell level using a 3D embryo culture system (Xiang et al., 2020). We sought to verify the GATA3^+^ ExE cells using these embryonic data as references. The expression dynamics of the lineage-specific genes that were defined as DEGs among three major lineages residing within d.p.f. 7-9 embryos, epiblast (EPI), primitive endoderm (PrE), and trophoblast (TrB), indicated an obvious trend of TrB gene induction in parallel with EPI gene downregulation (Figure S2D). For a more detailed comparison, we obtained the gene expression profile from the single cells treated with AP for 4 days, the time point when almost all of the cells became positive for GATA3. A total of 2,464 cells were divided into five clusters and two differentiation trajectories were estimated (Figures 2D-E). Profiling all clusters by DEGs among six trophoblast subpopulations of 6-14 d.p.f embryos revealed that cluster 2 shared high-expression genes with early-syncytiotrophoblasts (E-STBs) of embryos (Figure S2E). Consistent with this, signature scores for E-STBs were higher in cluster 2 than in the other clusters (Figures 2F-G). The protein expression of cluster 2-enriched genes, including the established markers for STBs (SDC1 and TBX3, Okae et al., 2018; Lv et al., 2019), was confirmed in a small proportion (Figures 2H-I, S2F). Re-analyses of bulk RNA-sequencing data using these marker sets indicated that expression of E-STB markers increased with the extension of the culture period (Figure 2J). The expression of HLA-G, a typical marker for extravillous trophoblasts (EVT), was detected, but the global trend of EVT-enriched gene expression did not support a robust ETV induction (Figures S2G-H). These results suggested that this culture condition was prone to induce biased differentiation to cells that were akin to STBs, rather than EVTs, of the peri-implantation embryo.

### Derivation of trophoblast stem-like cells from pESCs

The GATA3^+^ ExE cells could be maintained for a further extended period, but after day 8, they showed reduced proliferation and spontaneous detachment from the culture substrate, suggesting that the optimization of culture conditions is needed for prolonged culture. Recently, Okae et al. reported the derivation of self-renewing trophoblast cell lines (trophoblast stem cells (TSCs)) from villous cytotrophoblasts (vCTB) of the human first-trimester placenta (Okae et al., 2018). Using a similar approach, several groups have succeeded in deriving TSCs from human pluripotent stem cells and reprogramming intermediates from fibroblasts (Castel et al., 2020; Cinkornpumin et al., 2020; Dong et al, 2020.; Guo et al., 2021; Io et al., 2021; Liu et al., 2020; Mischler et al., 2021). We applied this method to GATA3^+^ ExE cells. The cells treated with AP for 4 days were harvested and transferred to the same culture conditions for TSC derivation. After a few passages, proliferative cells were obtained without any clonal selection (Figure 3A). These cells were expanded over several months (over 40 passages), retaining uniform expression of GATA3 as well as several trophoblast markers (Figures 3A-C). We call these cells trophoblast stem-like cells (TSLCs). To comprehensively evaluate the cell state, global gene expression of pESCs, GATA3^+^ ExE cells and TSLCs (after 10 or 15 passages) were profiled by mRNA-sequencing. A principal component analysis demonstrated that TSLCs with different passage numbers were clustered within a close region, indicating their ability to self-renew (Figure 3D). Otherwise, TSLCs were separated from both pESCs and GATA3^+^ ExE cells (Figure 3D). Consistent with these observations, the global expression level of the vCTB-associated genes was higher in TSLCs than in parental AP-treated cells, while STB-related genes identified above were lower (Figures S3A-B). It is noteworthy that the expression of some vCTB-associated molecules, including p63, a key transcription factor for vCTB stemness (Lee et al., 2007; Li et al., 2013), was remained low during all periods of AP-induced differentiation but upregulated after TSLC derivation (Figures 3E-F, S3C-D). Principal component analyses combined with published datasets (Okae et al., 2018) revealed that our TSLCs were closely clustered with vCTB-derived TSCs (Figure 3G). Moreover, the Spearman correlation test also indicated high similarity between TSLCs with reported TSC lines derived from vCTBs as well as naïve ESCs or reprogramed fibroblasts (Figures 3H, S3E-G). These observations indicated that TSLCs were different from AP-induced GATA3^+^ ExE cells, but analogous to TSCs.

**Figure 3.**
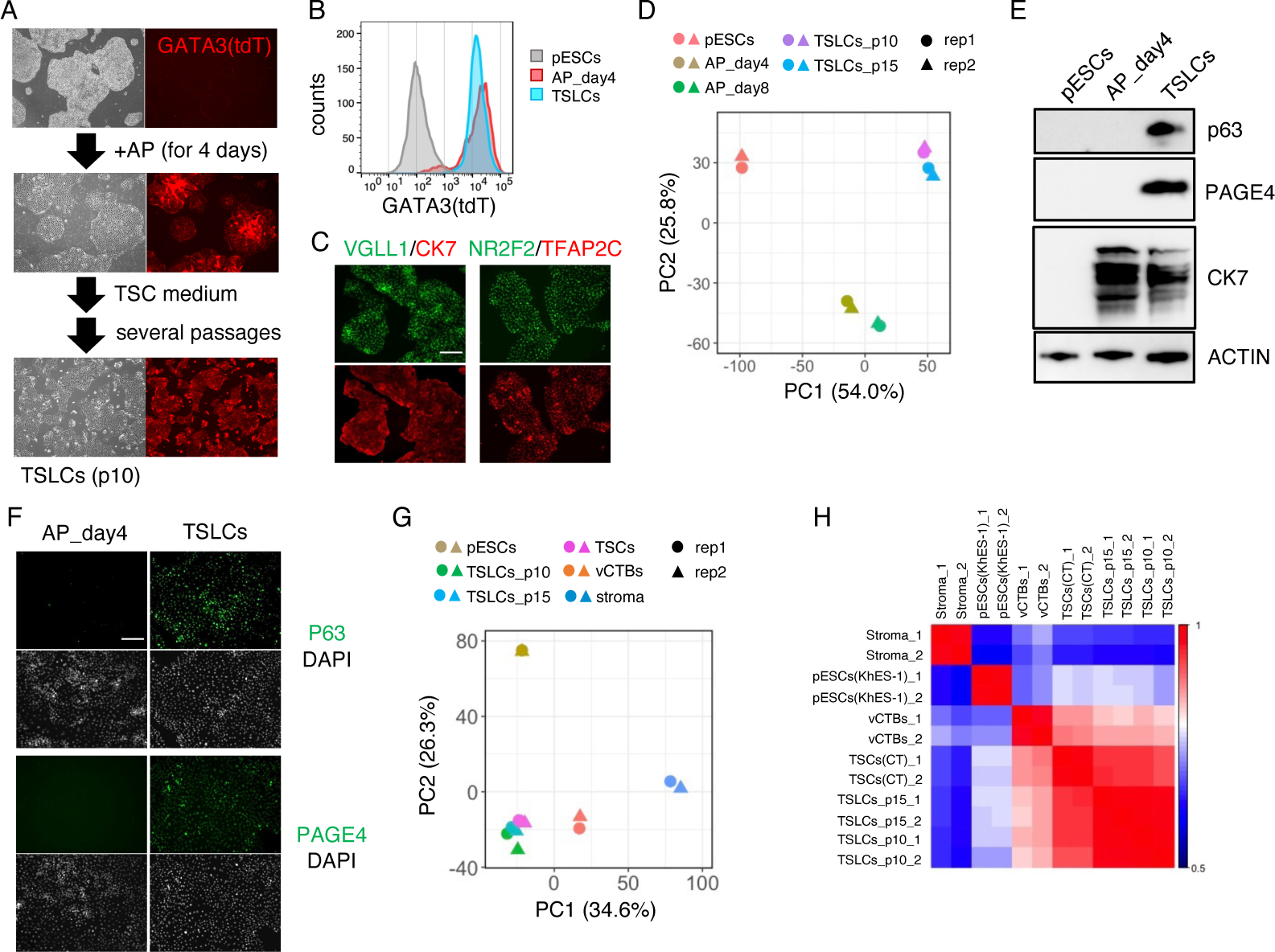
Derivation of trophoblast stem-like cells from pESCs. (A) Schematic representation of TSLC derivation from GATA3-knockin pESCs. Red fluorescence indicates the GATA3-expressing cells. (B) Flow cytometric panels showing GATA3 expression level in pESCs, differentiating cells (day 4) and established TSLCs. (C) Immunostaining using GATA3-knockin pESCs for pan-trophoblast markers. Scale bars show 100 µm. (D) Western blotting analyses for the indicated proteins in pESCs, AP-treated cells (day 4) and TSLCs. ACTIN was examined as a loading control. (E) Immunostaining using GATA3-knockin pESCs for the indicated TSLC markers. (F) PCA analyses of pESCs, AP-induced differentiated cells (days 4 and 8) and TLSCs (two passage numbers). The top 1,000 valuable genes were used for the variation calculations. (G) PCA analyses of pESCs, TLSCs (this study), TSCs, vCTBs, and placental stroma cells (Okae et al., 2018). The top 1,000 valuable genes were used for the variation calculations. (H) Spearman correlation of transcriptomes from pESCs and TSLCs with published datasets from TSCs and placental stroma cells. Scale bars show 200 µm.

All results described above support the idea that human pESCs have the competence to differentiate into trophoblast-like cells despite their primed state of pluripotency. It is interesting that inhibitor-mediated perturbation of pluripotent signals was sufficient to stimulate pESCs not only to exit from their undifferentiated state but also to undergo a sequential transition into GATA3^+^ cells. This suggests uncharacterized intrinsic programs for trophoblast-like fate conversion.

### Molecular analyses of pESC-to-GATA3^+^ ExE conversion

We sought to decipher the intrinsic mode of the differentiation program. The scheduled medium-switching study demonstrated that the first 3 days of treatment with the inhibitors was sufficient to produce GATA3-positive cells with over 90% efficiency (Figure S4A), indicating that this is a critical period for the fate determination of GATA3^+^ ExE cells. We examined BMP responses during this period, because the importance of BMP signaling to direct pESCs to trophoblasts has been repeatedly reported in the past decades (Roberts, 2018; Xu et al., 2002). Through a series of biochemical assays, we proved that BMP-SMAD signaling was transiently activated by AP stimulation (Figures 4A-D). When surveying RNA-sequencing data, we noticed that among BMP ligands, the mRNA expression of BMP4 was upregulated (Figures 4C, S4B). The secretion of BMP4 into the culture medium was confirmed (Figure 4D). Two different types of BMP signal blockers, NOGGIN (NOG) as an antagonist for extracellular BMPs and LDN-193189 (LDN) as a kinase inhibitor for intracellular signal cascade, impaired BMP-dependent transcription, ensuring the cell-autonomous activation of BMP signaling in the differentiating cells (Figure 4E). When BMP signal input was repressed by the combined action of NOG and LDN, the AP-induced GATA3-expressing cells were significantly decreased (Figures 4F-G, S4C-D). Alternatively, PAX6^+^ ectodermal cells were induced under these conditions (Figure 4G). Consistent with these results, quantitative RT-PCR analyses showed a reduction in trophoblast genes and the upregulation of ectoderm genes in the BMP-inhibited condition (Figure S4E). Few cells seemed to still express pluripotency-related factors (Figure 4G, S4E). These data support the hypothesis that upon exit from an undifferentiated state, cells begin to express BMP4, and this endogenous signal determines the fate of GATA3^+^ ExE cells while preventing ectodermal fate.

**Figure 4.**
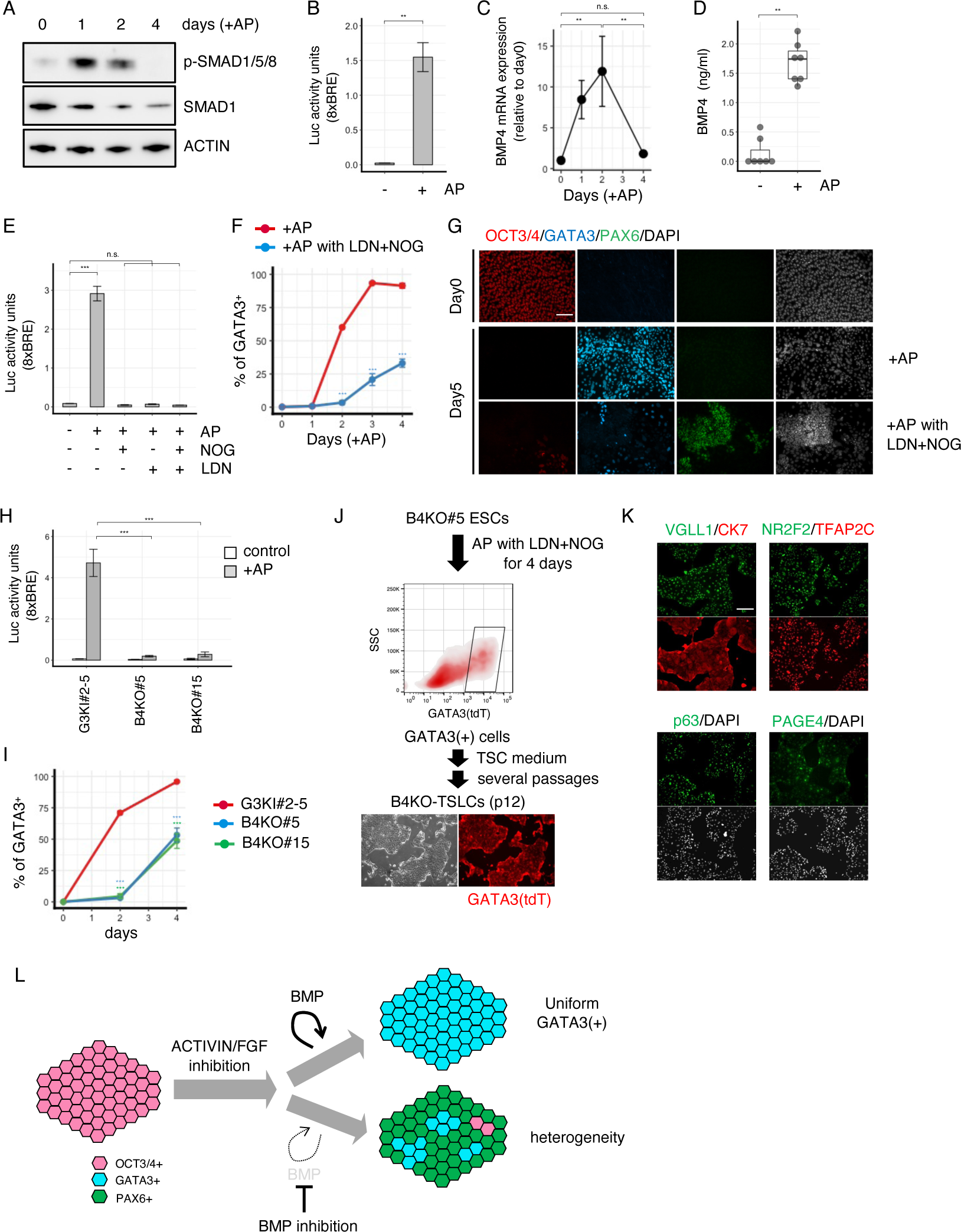
Molecular analyses of pESC-to-GATA3^+^ ExE conversion. (A) Western blotting analyses for the SMAD signaling pathway at the indicated timepoints of AP-induced differentiation. ACTIN was examined as a loading control. (B) Luciferase activity measurement using the 8×BRE reporter plasmid before and after AP addition. (C) qPCR assay for BMP4. Cells were treated with AP for the indicated periods. (D) Secretion levels of BMP4 in culture supernatants (day 3). (E) Inhibitory effects of NOG and LDN on BMP-dependent transcription. Luciferase activity was measured before and after AP addition (day 2). (F) Time course quantitation of GATA3^+^ cells in the presence and absence of BMP blockers. (G) Immunostaining using KhES-1 cells for the indicated lineage markers. (H) The impaired BMP responses upon BMP4 gene disruption. Reporter plasmids were transfected into G3KI#2-5 cells and two BMP4 knockout clones, and the activities of luciferase were measured before and after AP addition (day 2). (I) Time course quantitation of GATA3 expression dynamics using GATA3-knockin and two BMP4-deficient clones. (J) Schematic representation of TSLC derivation from a BMP4-deficient clone. (K) Immunostaining of B4KO pESC-derived TSLCs for the indicated markers. (L) Schematic diagram of BMP-dependent and -independent differentiation. Scale bars show 200 µm. Error bars in the graphs represent standard deviations. Paired t-test (n = 3) (B, E); Tukey’s test (n = 3) among all groups (C); Wilcoxon signed-rank test for paired data (D); unpaired Student’s t-test (n = 3) versus control cells at each time point (F, I); Durnett’ test (n = 3) versus control (day 0) (G); N. D., not detected, n. s., not significant; * *p* <0.05, ** *p* <0.01, *** *p* < 0.001.

We were interested in the observation that even in the presence of BMP blockers, approximately 30% of cells become GATA3^+^ cells, which also express other markers for trophoblasts (Figures 4F-G, S4D-E). This suggests that BMP inhibition was efficient but insufficient to deprive pESCs of trophoblast-like differentiation competence. To prove this possibility, we attempted to disrupt cell-autonomous BMP signal activation by genetic manipulation. We generated ESC lines in which exon 3 of the BMP4 gene was deleted (B4KO, clone #5 and #15) (Figures S4F-G). When these cells were treated with AP, BMP reporter activity was dropped nearly to basal levels, indicating functional impairment of BMP signals (Figure 4H). This approach rendered pESCs resistant to differentiation into GATA3^+^ cells, but a portion of cells still went toward this fate, as seen in the inhibitor experiments (Figures 4I, S4H). When the B4KO line was treated with AP in the presence of NOG/LDN, a condition in which endogenous BMP signal inputs were strongly depleted, GATA3^+^ cells still emerged (Figure 4J). By transferring these cells to the TSC medium, we succeeded in obtaining TSLCs (Figures 4J-K). These results demonstrated a BMP-independent pathway toward GATA3^+^ ExE cells, although cell-autonomous BMP signal activation is important for efficient conversion (Figure 4L).

### Identification of ELF3 as a determinant of extraembryonic fates

The BMP-independent differentiation into GATA3^+^ cells is intriguing but poorly characterized. To gain insight into the undescribed route, we searched for AP-induced genes that were upregulated both in the presence and absence of BMP blockers (Figure 5A). Among the commonly upregulated genes, which include genes encoding nine transcription factors, we found that an ETS-like transcription factor ELF3 induced GATA3^+^ cells (see below). Quantitative RT-PCR, western blotting, and immunostaining showed elevated expression of ELF3 in the early phase of AP-induced differentiation (Figures 5B-D). To examine the effect of ELF3 on fate specification, we generated cell lines in which HA-tagged ELF3 was induced by doxycycline (dox) (tet-E3, clone #3 and #5). In these cells, dox supplementation induced ELF3 protein expression within 24 h, and they were localized to the nucleus in all cells observed (Figures 5E-F, S5A-B). Even though we did not use any factors other than dox, these cells started to change their morphology and became positive for GATA3 within 48 h (Figures 5F-H, S5C). The emergence kinetics of GATA3^+^ cells seemed to be accelerated as compared to that of AP-induced differentiation (Figure S5D). These GATA3^+^ cells also expressed other trophoblast markers including placental hormone CGβ and placenta-specific ERVs (Figures 5H-K, S5G). By transferring these cells to the TSC media, we succeeded in obtaining TSLCs (Figures S5E-F). Similar to the AP-induced differentiation, ELF3 induced differentiation into the E-STB marker-positive cells (Figure S5G). The pulsed dox supplementation studies indicated that transient exposure to dox was sufficient for fate conversion (Figure S5H). The ineffective suppression of ELF3-induced differentiation by BMP blockers supported the expected BMP-independent action of ELF3 in this process (Figure 5G). To further validate the ELF3 function, we generated ELF3-deficient cell lines in which exon2-7 of the ELF3 gene were excised (E3KO, clones #21, #28) (Figures 5L, S5I-J). Although this genetic manipulation reduced the efficiency of the AP-induced differentiation, these cells still maintained their competency (Figure S5K). However, under conditions in which BMP activity was depleted, ELF3-null cells exhibited severe deficiency in generating GATA3^+^ cells upon AP treatment (Figure 5M). These observations demonstrate that ELF3 acts as a driver of the BMP-independent differentiation, and the cooperative action of BMP-dependent and -independent mechanisms to achieve efficient induction of GATA3-expressing cells from human pESCs.

**Figure 5.**
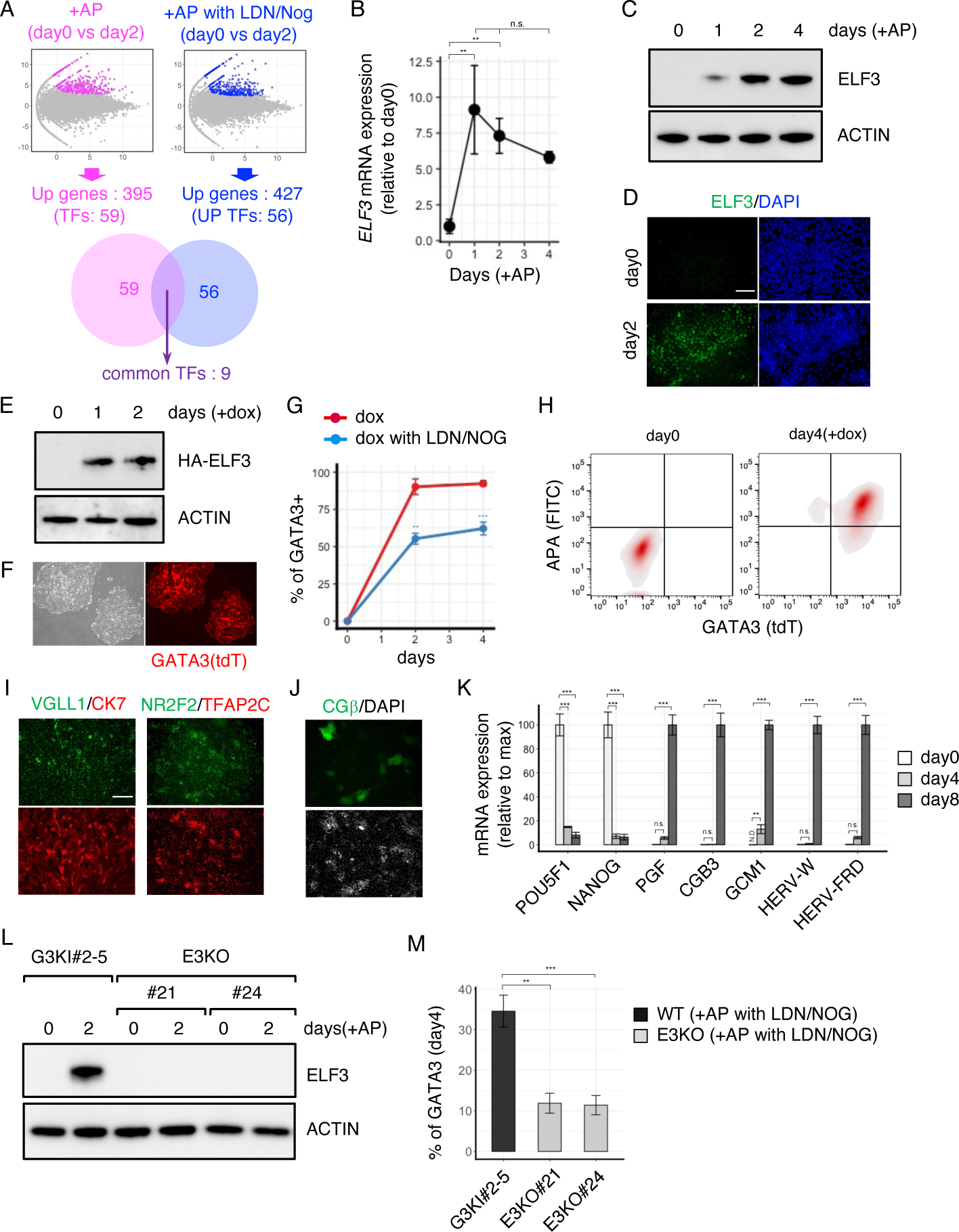
Identification of ELF3 as a determinant of extraembryonic fates. (A) Transcriptome comparison for the identification of BMP-independent upregulated genes. (B) qPCR assay for ELF3. Cells were treated with AP for the indicated periods. (C-D) Western blotting (C) and immunostaining analyses (D) showing the expression of ELF3 at early phase of differentiation. ACTIN was examined as a loading control. (E) Western blotting for dox-dependent induction of ELF3. ACTIN was examined as a loading control. (F) Induction of GATA3^+^ cells from tet-E3#3 cells after 4 days of dox addition. Red fluorescence shows GATA3-expressing cells. (G) Time course quantitation of dox-induced GATA3^+^ cells in the indicated time points (red line). GATA3^+^ cells were also quantified in the presence of BMP blockers (blue line). (H) Flowcytometric panels of dox-treated tet-E3#3 cells for APA^+^/GATA3^+^ cells at day 4 after dox addition. (I-J) Immunostaining for the indicated markers at day 4 (I) and day 6 (J) of differentiation. (K) qPCR assay for pESC and trophoblast genes before and after ELF3 induction. The tet-E3#3 cells were treated with dox for the indicated periods. (L) Western blotting for examining ELF3 expression in two ELF3 knockout clones and the parental cells. ACTIN was examined as a loading control. (M) Comparison of induction efficiency of GATA3^+^ cells in the presence of BMP blockers between two ELF3 knockout clones and the parental cells. Scale bars show 200 µm. Error bars in the graphs represent standard deviations. Tukey’s test (n = 3) among all groups (B); unpaired Student’s t-test (n = 3) versus control cells at each time point (G, M); Durnett’ test (n = 3) versus control (day 0) (K).

### Amnionic features at the early phase of differentiation

As described previously in this manuscript, the BMP4-expressing cells are key for determining the fate of GATA3^+^ ExE cells. Unbiased hierarchy clustering indicated that pESCs and AP-treated cells were classified into the same group until day 2, a time point when differentiating cells did not express GATA3 but expressed and secreted BMP4 (Figure S2A, 4C-D). We were interested in these observations because in primate peri-implantation embryos, BMP4 expression was predominantly observed in the nascent amnion (AM), an EPI-derived epithelium that resides beneath the trophoblast layer to form an amniotic cavity (Sasaki et al., 2016). Therefore, we tested the hypothesis that AP-treated cells in the early phase contained AM-like cells. Morphological changes implying the transition from columnar to squamous epithelium, an anatomical feature distinguishing AM from EPI (Figure S6A; Luckett et al., 1976; Xiang et al., 2020), were observed during AP-induced differentiation (Figures 6A-C). The squamous epithelium was negative for NANOG, but co-expressed OCT3/4 and TFAP2A at a low level (Figure 6D), as seen in AM of primate embryos or stem cell-based models for human AM (Shao et al., 2017; Xiang et al., 2020; Zheng et al., 2019; Zhu et al., 2020).

**Figure 6.**
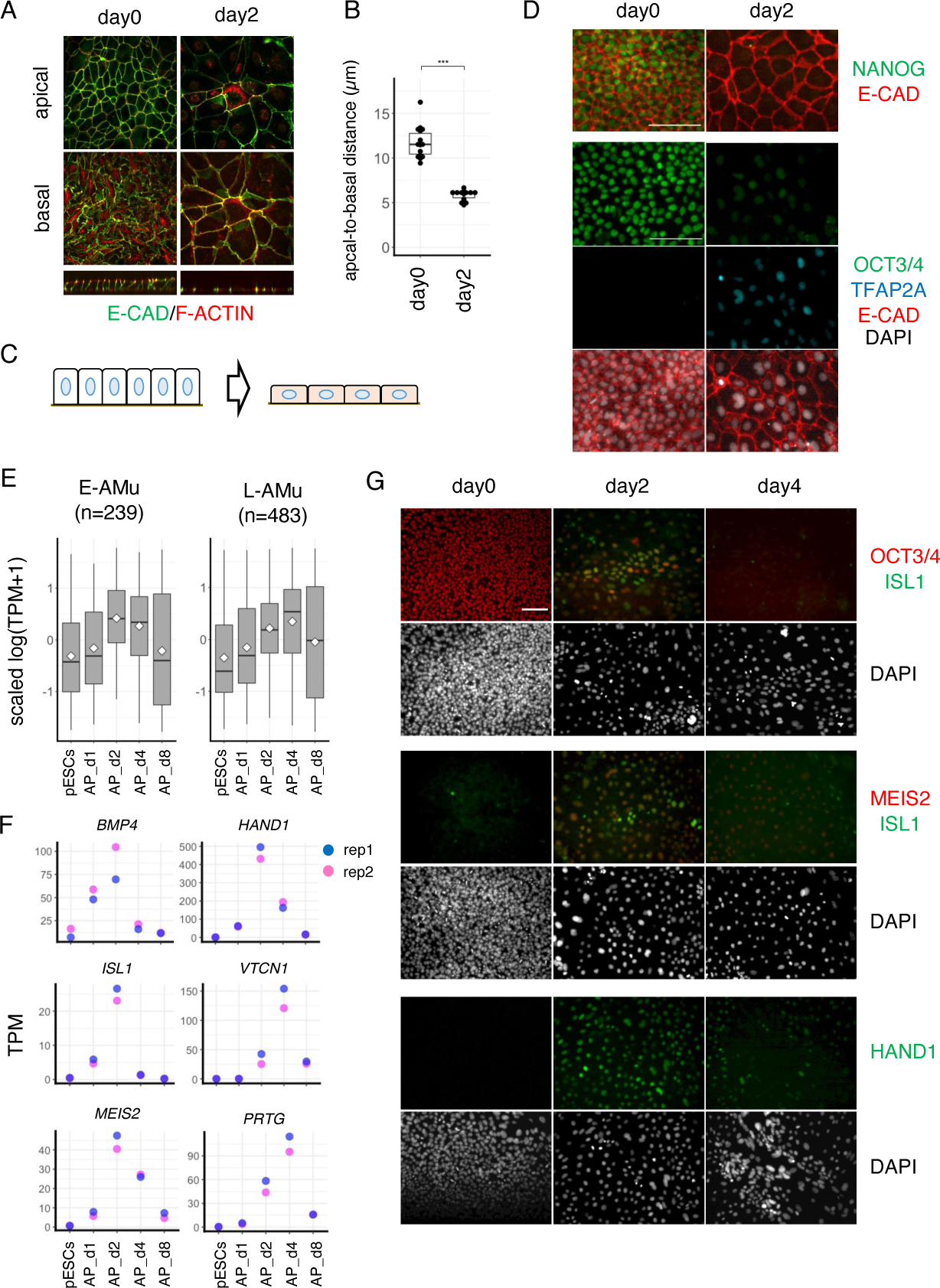
Amnionic features at the early phase of differentiation. (A) Confocal images of the KhES-1 colonies treated or untreated with AP for 2 days. The upper and middle panels are the x-y views at apical and basal sections, respectively. The low is the y-z view. (B) Comparison of apical-basal distances of AP-treated or -untreated colonies (n = 10). (C) Schematic diagram of the morphological transition (y-z view) of the AP-treated colony. (D) Immunostaining of KhES-1 cells treated with AP for the indicated periods. (E) Expression dynamics of the selected AM genes during AP-induced differentiation. The scaled TPM values for E-AMu (n= 238) and L-AMu genes (n = 483) are shown. (F) Dot plot presentation showing the expression dynamics of the early and late AM marker genes based on bulk-mRNA-sequencing data. (G) Immunostaining of AP-treated KhES-1 cells for the AM-related proteins at the indicated timepoints of differentiation.

AM segregation from post-implantation EPI before gastrulation is unique to primates (Enders et al., 1986). Several technical and ethical issues have hampered the understanding of human AM properties at the molecular level, but the transcriptional feature of cynomolgus monkey AM is available at single-cell resolution (Ma et al., 2019; Niu et al., 2019). Since the AM genes, which are classified as DEGs relative to EPI, overlap with typical trophoblast genes, we generated AM-unique gene list by subtracting placenta-enriched genes from the reported AM gene list (Figure S6B) (Ma et al, 2019; Cao et al, 2020; see Methods). We first examined the temporal expression of the early and late AM-unique genes and found that they showed transient upregulation around days 2-4 of AP-induced differentiation (Figures 6E-F, S6C). After selecting some genes, the expression of which was predominantly abundant in human AM (Figure S6D), we evaluated their protein expression during AP-induced differentiation by immunostaining. These analyses confirmed that AM-associated markers, such as ISL1 and HAND1 (Knöfler et al., 2002; Ma et al., 2020; Yang et al., 2020), were induced and co-expressed at day 2 (Figure 6G). Although their expression levels seemed to be relatively low, these results support our speculation that AM-like cells appeared in the early stages of differentiation.

### Bifurcation of differentiation trajectory into amnion- and syncytiotrophoblast-like cells

The expression of BMP4 and ISL1 returned to the basal level immediately, but some AM markers were still expressed at day 4, a time point when E-STB-like cells appeared (Figures 6F-G). We again analyzed the single-cell transcriptome dataset to define which cells expressed them, and found their biased distribution to cluster 1 and 4 populations (Figure 7A). Scoring gene expression signature using the late AM-unique geneset revealed that cluster 1/4 showed a higher score than other clusters, including the aforementioned E-STB-like population (cluster 2) (Figures 7B-C). By comparing gene expression between clusters 1 and 2, we noticed that the cluster 1 cells highly expressed several genes that had been reported to be specific for the first-trimester human amnion (Roost et al., 2015; Zhu et al., 2020) (Figures S7A-B). Among them, cytokeratin proteins CK17 and CK23 were identified to be useful for distinguishing them (Regauer et al., 1985; Turco et al., 2018) (Figures 7D, S7C). Flow cytometric analyses indicated that CK17^+^ and CK23^+^ populations simultaneously emerged around day 4 in a mutually exclusive manner (Figure 7E). Immunostaining analyses revealed the spatial compartmentation of these populations; CK23 signals were clustered in the inner regions and frequently overlapped with the STB marker SDC1, while CK17 was expressed as a sporadic pattern and co-localized with HAND1 (Figures 7F, S7D). In line with the mRNA expression, the AM-related molecules were rarely expressed in CK23^+^ cells, and conversely TBX3 or SDC1 was absent in the CK17^+^ cells (Figure S7E). Syncytialization and CGβ expression, two salient phenotypes witnessed in embryonic STBs, were observed exclusively in CK23-positive cells, and never overlapped with HAND1/CK17 signals (Figures 7G-H). The pESC-derived syncytia exhibited striking immunological hallmarks; they expressed immuno-suppressive PD-L1 while expressing a negligible level of class I HLA molecules (Figures 7I-L, S7F). This phenotype might be implicated in their evasive potential from the maternal immune system. Taken together, we concluded that the STB-like cells stemmed from the AM differentiation trajectory (Figure 7M). While STB-related genes continued to increase their expression, AM-related genes showed a tendency to decline with prolonged culture (Figures S7G-H), implying that the culture condition was inferior for further differentiation to the mature amnion membrane.

**Figure 7.**
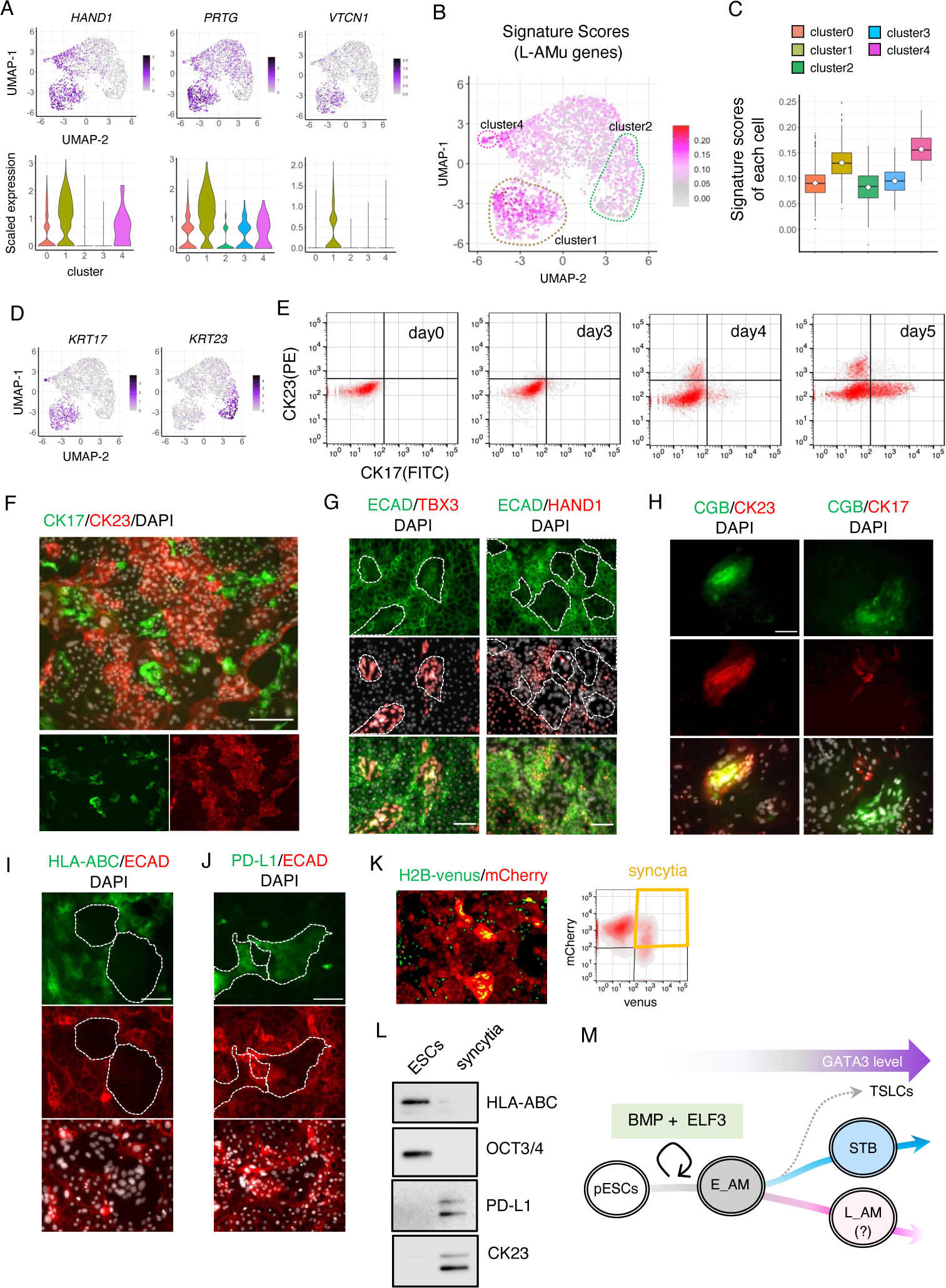
Bifurcation of differentiation trajectory into amnion- and syncytiotrophoblast- like cells. (A) Violin and scatter plot representations of the selected AM genes. Cell clusters were the same as those in Figure 2D. (B-C) Signature score distribution at day4 of differentiation. Signature scores were calculated using genesets for the late AM (n = 494) and applied onto a UMAP space (B). The score in each cluster are presented as a boxplot (C). (D) Scatter plot representation of cytokeratin 17 or 23 expression in each cell. (E) Flow cytometric panels for CK17/CK23 expression. The KhES-1 cells were treated with AP for the indicated periods. (F-H) Immunostaining of KhES-1 cells treated with AP for the indicated periods. Co-staining for cytokerain-17 and -23 (F, day 5), ECAD and TBX3 or HAND1 (G, day 6), CGβ and CK23 or CK17 (H, day 6). (I) Immunostaining of HLA-ABC in KhES-1 cells treated with AP for 8 days. Syncytium were defined with E-CAD signal, and indicated as the enclosed regions by dotted lines. (J) Immunostaining of PD-L1 of KhES-1 cells treated with AP for 8 days. PD-L1-pisitive areas are indicated as enclosed regions (dotted lines). (K) Isolation of syncytial cells from the differentiated culture (at day 8). Syncytial cells are defined by dual fluorescence expression as described Figure 1K (left image), and indicated by an orange polygon (in the flow cytometric panel, right). (L) Western blotting analyses to evaluate the level of expression of HLA-ABC and PD-L1 in the undifferentiated pESCs (lane 1) and purified syncytial cells (lane 2). The same cell number of each sample was collected by flow cytometry, from which cell lysate was generated. (M) Schematic model for STB branching from AM differentiation trajectory. Scale bars show 100 µm (I, J) or 200 µm (F, G, H).

## Discussion

Human ESCs are known to readily acquire certain trophoblast characteristics in response to BMP ligands. However, trophoblast identity of the ESC derivatives is often suspected, because conventional human ESCs (pESCs) have been regarded to recapitulate the restricted differentiation competence of the post-implantation epiblasts. Some reports have claimed that they represent non-trophoblastic cells, such as mesodermal or amnionic derivatives (Bernardo et al., 2011; Guo et al., 2021; Ito et al., 2021). Considering the controversial situation, we avoid referring the AP-induced cells as ‘trophoblasts’ but alternatively as ‘GATA3^+^ ExE cells’, and characterized them exhaustively. We demonstrated that pESCs differentiate into hormone-producing syncytia, analogous to the early STBs of post-implantation human embryos. The successes in TSLC derivation imply the presence of a progenitor-type population within this culture. Some CTB-associated genes are expressed at a low or undetectable level during AP-induced differentiation, but these genes were shown to be upregulated after TSLC derivation. Although it is ambiguous whether the STB-like syncytia are derived through the progenitor state corresponding to embryonic CTBs, these results indicate that human pESCs possess the competence to give rise to not only STBs in primitive syncytium of the implantation embryo but also an *in vitro* counterpart of villous CTBs in the first-trimester placenta. In a previous report, monkey pESCs were shown to contribute to the placental tissue when injected into the morula stage of the monkey embryo, supporting our results (Kang et al., 2018).

A key finding of this study is that exogenous BMP ligands are not essential for GATA3^+^ ExE fate specification from pESCs. It is widely known that mimicking signal communication between embryonic and extraembryonic tissues directs pluripotent cells to initiate differentiation into mesendodermal or germ lineages, but our results suggest that human pESCs exert their differentiation potency into AM-to-STBs without the help of any external signals. Taking advantage of this culture condition, we uncovered two intrinsic genetic programs that steer differentiating cells into GATA3-expressing lineages. Upon exit from the undifferentiated state, cells begin to express and secrete BMP4, which then initiates the well-studied trophoblast-like differentiation. Thus, BMP-mediated intercellular communication governs cellular behaviors to allow uniform conversion to GATA3^+^ cells. When BMP activity is eliminated from the culture, differentiating cells may stochastically select their directions. Although ectodermal fate became predominant in the given situation, some cells still chose extraembryonic fate. We identified the transcription factor ELF3 as another determinant of extraembryonic fate specification. ELF3 is induced independent of BMP signals, and can initiate BMP-independent differentiation to GATA3^+^ cells without any other additives. This differentiation competence is deprived by the combined impairment of BMP4-mediated intercellular communication and ELF3-mediated cell-intrinsic program, highlighting the cooperation of these two distinct modes of genetic code to guide epiblast-like stem cells to GATA3^+^ lineages.

The pESC-derived GATA3^+^ cells express many genes implicated in placental development and function and give rise to cells that resemble STBs in peri-implantation embryos. From the viewpoint of developmental lineage, however, they are likely to be distinct from conventional trophoblasts. In our differentiation system, we recognized nascent AM-like cells as a transient cell state, which partially agree with the previous reports (Guo et al., 2021; Io et al., 2021). They express definitive AM markers, including BMP4 and ISL1, and are functionally crucial for the subsequent steps of differentiation because the inhibition of the BMP4 action impairs the increase of GATA3^+^ cells. The expression of some AM genes is peaked around days 2-4 of differentiation, after which STB-like cells emerged. We found that a proportion of cells were still positive for certain AM-associated genes at day 4, but interestingly, they were clustered as a distinct population from the cells undergoing STB differentiation. Thus, the GATA3^+^ ExE cells are the intermingled pool of amnion-like cells and STB-like cells, suggesting a bifurcated trajectory of AP-induced differentiation, maturation to future amnion membrane and placental hormone-producing syncytial cells (Figure 7I). We supposed that the AP-induced STB-like cells are phenotypically equivalent to the BAP-induced ones (Xu et al., 2002; Yabe et al., 2016), but the elimination of exogenous inducers from the culture conditions sheds light on the critical role of BMP4-expressing nascent AM-like state linking pESCs to STB-like cells.

Finally, we discuss the biological implications of differentiation of STB-like cells from epiblast-like pESCs. Whether ESC-derived trophoblast-like cells are truly *in vitro* counterparts of authentic embryonic trophoblasts has been questioned since the first report by Thompson’s group (Xu et al., 2002). However, recent progress in primate embryology has unveiled substantial differences between mouse and human development (Nakamura et al., 2016; Petropoulos et al., 2016; Rossant and Tam, 2017; Turco and Moffett, 2019), and offers another explanation for this long-standing controversy. The primate amnion has been shown to express several typical markers of trophoblasts, despite the high similarity to epiblast in the global transcriptome (Ma et al., 2020; Niu et al., 2020; Xiang et al., 2020). Considering together with these observations, our results raise the possibility that the amnion provides STB-like endocrine syncytia. This might represent an alternative differentiation to hormone-producing syncytia that has been overlooked in rodent embryogenesis. It is a delicate issue which would be better to refer to the amnion-derived STB-like cells as ‘amnion’ or ‘trophoblast’, but the pESC-derived ones shared fundamental properties with the embryonic STBs in gene expression signature, endocrine functions, immunological features and characteristic syncytial morphology. Based on these observations, we propose the following hypothesis: the ‘secondary’ STB specification from human nascent amnion at the post-implantation stage. They may back up the trophectoderm-derived primitive STBs to support the growth of the human fetus which is entirely embedded into the maternal endometrium. Although the presence of STB-like syncytia within the amnion epithelium remains unclear, it has been reported that CGβ-expressing cells are located proximal to the amniotic epithelium of human embryos (Xiang et al., 2019). Whether these cells are derived from the blastocyst trophectoderm or amnion is an exciting question that challenges the traditional view of mammalian development. Future *in vivo* investigations utilizing strict lineage tracing techniques will help us to discuss the hypothesis that originates from *in vitro* studies.

## Acknowledgements

We thank K. Matsuura, H. Niwa, T. Yamamoto, T. Tsukiyama, M. Mutou, M. Ema, and all lab members for their technical support and useful discussions. We also express special thanks to Yoshiki Sasai with respect to his legacy in science. This work was supported by the Japan Agency for Medical Research and Development (17bm0704013h9902); Takeda Science Foundation; the Mochida Memorial Foundation for Medical and Pharmaceutical Research; Joint Usage/Research Center for Developmental Medicine, IMEG, Kumamoto University; ISHIZUE 2020 of Kyoto University Research Development Program, and INFRONT Office of Directors’ Research Grants Program.

## Author Contributions

M.O. designed the project, performed experiments and data analyses, and wrote the manuscript. M.E. supported M.O. in data analyses.

## Experimental procedures

### ESC culture

All experiments using the hESC lines (KhES-1 and KthES-11) were performed following the hESC guidelines of the Japanese government. The 253G1 hiPSCs were a gift from S. Yamanaka (Kyoto University). Undifferentiated hESCs/iPSCs were maintained as previously described (Ohgushi et al., 2015). Cells were cultured on feeder layers of mouse embryonic fibroblasts (MEF; Kitayama Labes; inactivated with 10 µg/ml mitomycin C and seeded at 4 × 10^5^ per 6 cm dish) in DMEM/F12/KSR medium (D-MEM/F12 (Sigma) supplemented with 20% KSR additive, 2 mM glutamine, 0.1 mM non-essential amino acids (Invitrogen) and 0.1 mM 2-mercaptoethanol). Recombinant human basic FGF (5 ng/ml, Wako) was added soon after seeding. For cell passaging, ESC colonies were detached by treating them with CTK dissociation solution at 37°C for 5-7 min, tapping the cultures, and then flushing them with a pipette, and recovering *en bloc* from the feeder layer. The detached ESC clumps were broken into smaller pieces by gently pipetting several times, and then these small clumps were transferred onto a MEF-seeded dish. For feeder-free cultures, contaminating MEF cells were removed by incubating the suspension on a gelatin-coated plate at 37°C for 2 h in the maintenance culture medium. In this procedure, MEF cells adhere to the dish’s bottom, but ESC clumps do not. The MEF-free hESC clumps were suspended in a MEF-conditioned medium and seeded onto a Matrigel substrate (BD Biosciences). The culture medium containing bFGF was refreshed daily until the next passage.

### Reagents and antibodies

Reagents were purchased from Tocris (Y-27632), Sigma (puromycine, G418), Wako (A83-01, PD03714 and LDN-193189), Clonthech (doxycycline), R&D (recombinant protein NOGGIN and BMP4), and Clontech (recombinant Cas9 proteins). ELISA kits were purchased from Abcam (human BMP-4 and CGβ) and Proteintech (human GDF15). Antibodies used in this work are shown in Table S1.

**Table S1.**
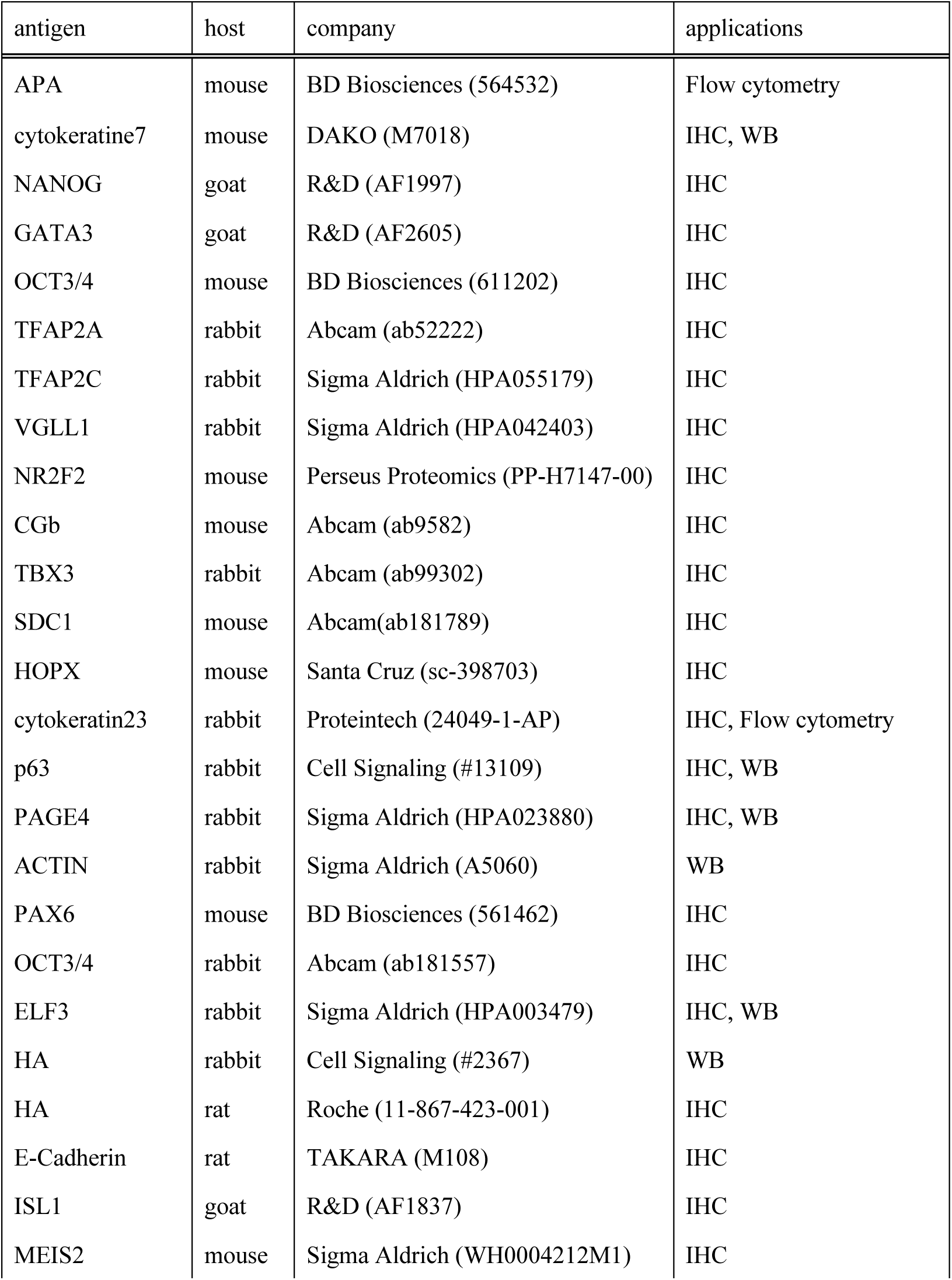
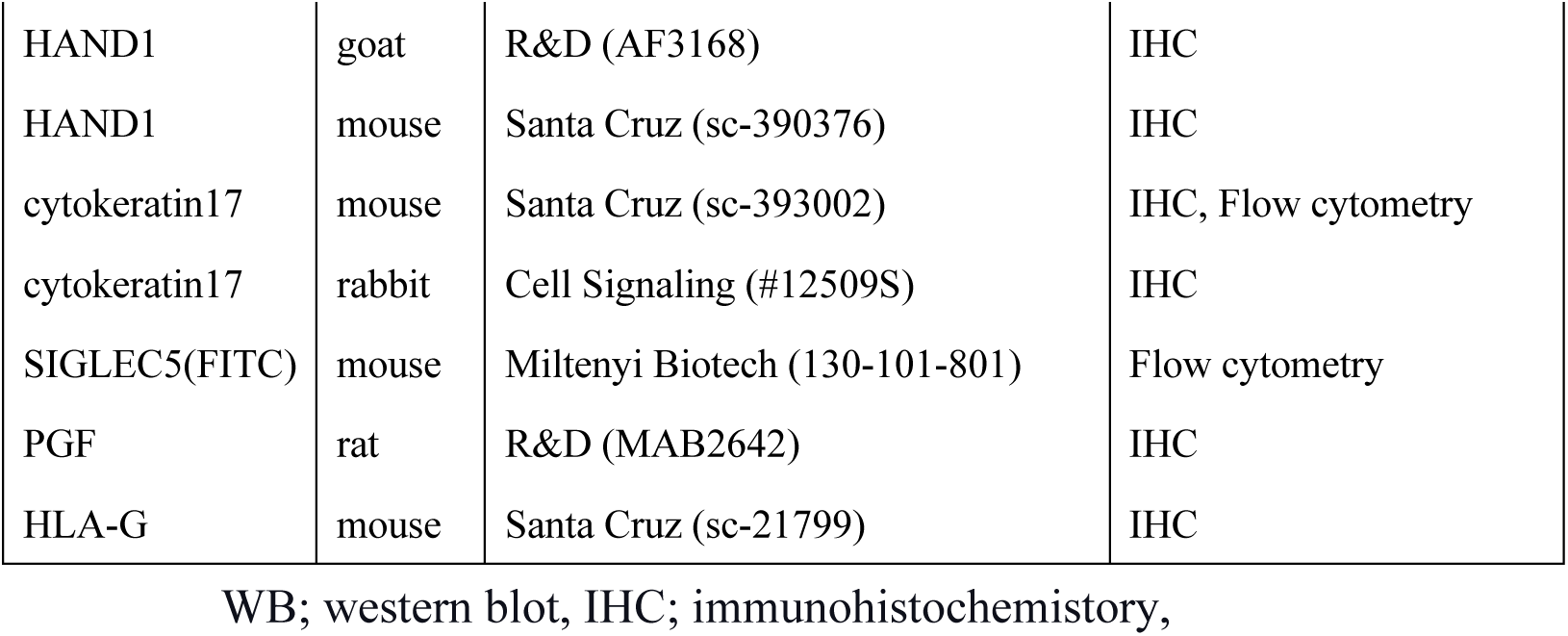
Antibodies used in this study.

| antigen | host | company | applications |
| --- | --- | --- | --- |
| APA | mouse | BD Biosciences (564532) | Flow cytometry |
| cytokeratine7 | mouse | DAKO (M7018) | IHC, WB |
| NANOG | goat | R&D (AF1997) | IHC |
| GATA3 | goat | R&D (AF2605) | IHC |
| OCT3/4 | mouse | BD Biosciences (611202) | IHC |
| TFAP2A | rabbit | Abcam (ab52222) | IHC |
| TFAP2C | rabbit | Sigma Aldrich (HPA055179) | IHC |
| VGLL1 | rabbit | Sigma Aldrich (HPA042403) | IHC |
| NR2F2 | mouse | Perseus Proteomics (PP-H7147-00) | IHC |
| CGb | mouse | Abcam (ab9582) | IHC |
| TBX3 | rabbit | Abcam (ab99302) | IHC |
| SDC1 | mouse | Abcam(ab181789) | IHC |
| HOPX | mouse | Santa Cruz (sc-398703) | IHC |
| cytokeratin23 | rabbit | Proteintech (24049-1-AP) | IHC, Flow cytometry |
| p63 | rabbit | Cell Signaling (#13109) | IHC, WB |
| PAGE4 | rabbit | Sigma Aldrich (HPA023880) | IHC, WB |
| ACTIN | rabbit | Sigma Aldrich (A5060) | WB |
| PAX6 | mouse | BD Biosciences (561462) | IHC |
| OCT3/4 | rabbit | Abcam (ab181557) | IHC |
| ELF3 | rabbit | Sigma Aldrich (HPA003479) | IHC, WB |
| HA | rabbit | Cell Signaling (#2367) | WB |
| HA | rat | Roche (11-867-423-001) | IHC |
| E-Cadherin | rat | TAKARA (M108) | IHC |
| ISL1 | goat | R&D (AF1837) | IHC |
| MEIS2 | mouse | Sigma Aldrich (WH0004212M1) | IHC |
| HAND1 | goat | R&D (AF3168) | IHC |
| HAND1 | mouse | Santa Cruz (sc-390376) | IHC |
| cytokeratin17 | mouse | Santa Cruz (sc-393002) | IHC, Flow cytometry |
| cytokeratin17 | rabbit | Cell Signaling (#12509S) | IHC |
| SIGLEC5(FITC) | mouse | Miltenyi Biotech (130-101-801) | Flow cytometry |
| PGF | rat | R&D (MAB2642) | IHC |
| HLA-G | mouse | Santa Cruz (sc-21799) | IHC |

WB; western blot, IHC; immunohistochemistry,

### Plasmids and transfection procedure

The cDNAs for HA-tagged ELF3 were amplified from the KhES-1 cDNA library by PCR using PrimeSTAR GXL (Takara) and were subcloned into the pENTR/D entry vector (Invitrogen). To achieve dox-dependent control of ELF3 expression, cDNAs were transferred into a piggyBac vector PB-CMV* by LR recombinase (Invitrogen). In this vector, cDNAs were driven by a minimal CMV promoter linked to a Tet-responsible element. To construct a reporter plasmid for BMP-dependent transcription, double-stranded oligonucleotides containing eight BMP-responsive elements in tandem (8×BRE; AGATCCTCTGGTCACAGGATAATAATCCTGACGCCAGAAAGTCTGGAGGTC) were synthesized (GeneArt, Invitrogen) and introduced into the pNL3.2[Nluc_minP] vector (Promega). All DNA fragments integrated into the vectors were sequenced using the appropriate primers. KhES-1 cells stably expressing fluorescence proteins were generated using a Piggybac (PB) transposon system, as described previously (Ohgushi et al., 2015). cDNAs for venus-tagged H2B and mCherry were subcloned into the PB transposon vectors containing a CAG-promoter-driven expression cassette, followed by an IRES-NeoR (a gift from Dr. Niwa). These PB vectors were co-transfected with a *pCAG-PBase* expression vector using the Lipofectamine Stem reagent (Invitrogen). A few days after the transfection, cells were passaged to the DR4 MEF (Cell Systems)-coated dishes and, on the following day, the medium was switched to a 100 µg/ml G418- containing one to obtain polyclonal stable pools with G418-resistance. Cells expressing fluorescence proteins were collected uisng a FACS Aria III flow cytometer (BD Biosciences), and used for downstream experiments.

### Generation of GATA3 reporter cell line

The gene-targeting strategy for G3KI ESC lines is illustrated in Figure S1A. The guide RNA was designed to target an immediate downstream of the stop codon of the human GATA3 gene and generated using the Guide-it sgRNA In Vitro Transcription System (Clontech). The target sequences of gRNAs are shown in Table S2. To create a donor template, homology arms to the integration site of the GATA3 gene, the 5′ arm (772bp) and 3′ arm (648 kbp), were amplified by PCR using the genome extracted from KhES-1 cells as a template. To eliminate the gRNA target in the 3’ arm of the donor, the PAM sequence was disrupted by PCR-mediated mutagenesis. The 5’ arm was linked with a cDNA for encoding P2A peptide-fused tandem tomato fluorescent proteins in flame. These fragments were integrated into a PL552 vector (Addgene, #68407) that contained a floxed expression cassette of puromycin-resistant genes downstream of the pgk promoter and sequenced. Using this vector as a template, a single-stranded donor for homologous recombination was generated using the Guide-it Long ssDNA Production System (Clontech). The gRNA, single-stranded donor, and recombinant Cas9 proteins (Clontech) were introduced together into the KhES-1 cells by electroporation (Neon device, Invitrogen). Recombinant ESCs were selected with 1 µg/ml puromycin, and the clones harboring both recombined and intact alleles were identified by genomic PCR and sequencing. The resultant clones were transfected with a Cre recombinase expressing vector, and subclones in which a pgk-PuroR cassette was removed were subjected to experiments.

**Table S2.**
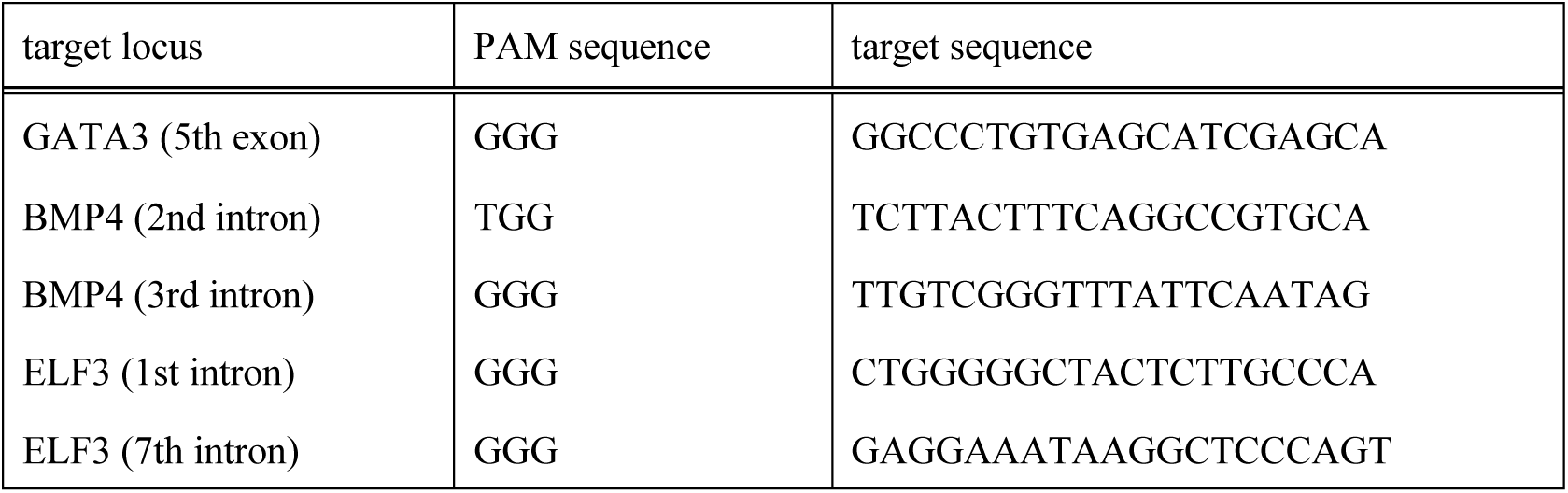
Target sequences for gRNA

### Generation of knock out cell lines

A set of gRNAs for each gene was designed as illustrated in Figure S4F (for BMP4) and S5G (for ELF3). The gRNA expression vectors were constructed using a Cas-9 sgRNA vector (Addgene, #68463) as a backbone. The target sequences of gRNAs are shown in Table S2. The expression vector for Cas9-2A-eGFP was obtained from Addgene (#44719). These vectors were co-transfected into GATA3 reporter cells (G3KI#5-2) using the Lipofectamine Stem reagent (Invitrogen). Two days after transfection, eGFP-positive cells were collected by flow cytometry and seeded onto MEF-feeders at a low cell density. The exon deletion was evaluated by genome PCR and sequencing, and clones lacking the targeted exons in both alleles were selected. The functional abolition of BMP4 was confirmed using a reporter assay. The lack of functional ELF3 protein was confirmed by western blotting.

### Generation of tet-E3 cell lines

GATA3 reporter lines (G3KI#5-2) were transfected with the *PB-CMV\** vector encoding *HA-tagged ELF3* together with the *piggyBac* vector pPB-CAG-rtTA-IRESneo. After treatment with G418, several drug-resistant clones were selected, and the clones harboring the transgene insertions were selected.

### GATA3^+^ ExE Differentiation

For GATA3^+^ ExE differentiation, human ESCs were collected as clumps, suspended in MEF-conditioned medium, and transferred onto Matrigel-coated dishes. The next day, the culture medium was replaced with fresh media containing A83-01 (1 µM) and PD173074 (0.1 µM), and the medium was refreshed daily. Unless otherwise indicated, fresh DMEM/F12/KSR without bFGF was used as the basal medium. We also tested other media such as MEF-conditioned DMEM/F12/KSR, mTeSR1(Stem Cell Technologies), StemFit AK-03N (AJINOMOTO), CDM (IMDM + Ham’s F-12 at 1:1 (Sigma), chemically defined lipid concentrate (Sigma), monothioglycerol (450 μM), 1% ITS-X supplement (WAKO) and purified BSA (3%, WAKO)) and N2B27 (DMEM/F12/GlutaMax + Neurobasal at 1 : 1 (Sigma), 1 × B27 (Gibco), 1 × N2 (Gibco) and 0.1 mM 2-mercaptoethanol) on the different substrates-coated plates (MEF-feeder cells, Matrigel and Laminin-E8 (Nippi)).

### TSLC derivation

ESCs (G3KI#2-5, B4KO#5 and tet-E3#3) were cultured in an AP-containing medium for 4 days. These cells were dissociated by TlypLE select, suspended in TSC medium (DMEM/F12 supplemented with 0.1mM 2-mercaptoethanol, 0.2% FBS, 0.3% BSA, 1% ITS-X supplement, 1.5 mg/ml L-ascorbic acid, 50 ng/ml EGF, 2 µM CHIR99021, 0.5 µM A83-01, 1 µM SB431542, 0.8 µM VPA and 5 µM Y27632), and seeded onto a collagen-VI-coated 6 well plate at 2.5 × 10^5^ cells per well. For the B4KO#3 clone, GATA3-positive cells were collected by flow cytometer and cultured under TSC derivation conditions. For TSLC derivation from tet-E3 clone #3, after treatment with dox for 2 days and cultured with dox-free media for another day, cells were dissociated and then transferred to the TSC derivation condition. The culture medium was replaced every two days. For cell passaging, cells were dissociated with TrypLE select and transferred to a collagen IV-coated plate. After a few passages, the cells started to show rapid proliferation while retaining a uniform GATA3 expression.

### Immunostaining

Immunostaining was performed as described previously (Ohgushi et al., 2015). The cells were fixed with 4% PFA at 4°C for 20 min and then permeabilized with 0.3% Triton- X100 solution. For HLA-ABC staining, cells were fixed and permeabilized with cold methanol at -20 °C for 10 min. After incubation in blocking solution such as 2% skim milk or 10% normal donkey serum (Abcam), cells were incubated in the blocking solution containing specific antibodies. The staining was visualized using secondary antibodies conjugated with Alexa Fluor-488, -546, or -647 (Invitrogen). For F-actin staining, Alexa Fluor-conjugated phalloidin (Invitrogen) was used. The nucleus was stained with DAPI or YOYO-1 (Invitrogen). Experiments were performed at least three times. Images were obtained using a fluorescence microscopy (LASX system, Zeiss) or an inverted microscope (IX81-ZDC, Olympus). For confocal observations, serial images were collected using a CSU-W1 unit (Yokogawa) configured with an IX81-ZDC microscope. Image processing was performed using the FIJI software.

### Western blot analyses

Cells were washed with PBS and treated on the plate with HEPES lysis buffer (50 mM HEPES pH 7.4, 150 mM NaCl, 1 mM EDTA, 1 % NP-40, and protease inhibitor cocktail) for 10 min at 4 °C with gentle shaking, and total cell extracts were harvested by pipetting. Immediately after adding the appropriate amount of 4 × LDS sample buffer (Invitrogen) to the extracts, the cells were subjected to a brief sonication for complete lysis. After boiling, ell lysates were analyzed by SDS-PAGE and western blotting. A 5 % skim milk solution was routinely used as a blocking reagent. In particular, for the detection of phosphorylated proteins, a 2 % BSA solution was used for blocking. Images were obtained with an Amersham Imager 600 (Fuji Film) and processed using the FIJI software.

### Flow cytometry analyses

To quantify fluorescence protein-expressing cells, cells were dissociated into single cells by TlypLE select and suspended in PBS containing the DNA dye DRAQ-7 (Cell Signaling). Using FACS Aria IIIu flowcytometry, the DRAQ-7-positive population was eliminated as dead cells, and the fluorescence in these individual cells was measured. For the detection of surface protein expression in live cells, dissociated cells were suspended in a culture medium and stood in suspension for 30 min in a CO_2_ incubator. The cells were washed with FACS buffer (PBS containing 2% fetal bovine serum) and incubated for 30 min in FACS buffer containing primary antibodies. When needed, staining with fluorescence-labeled secondary antibodies was performed. After washing and suspending in FACS buffer containing DRAQ-7, the cells were subjected to flow cytometry analyses. For cytokeratin staining, dissociated cells were fixed with chilled methanol for 10 min. The fixed cells were incubated in blocking solution (10% normal donkey serum) for 30 min, and then incubated in the blocking solution containing primary antibodies for 1 h (or overnight). The cells were stained using secondary antibodies conjugated with AlexaFluor-488 or -546 (Invitrogen), and subjected to flow cytometry analyses.

### Luciferase reporter assays

The 8 × BRE reporter plasmid was transfected into ESCs together with the pGL3.2 plasmid for normalization. Cell extracts were prepared using Passive Lysis Buffer (Promega), and luciferase reaction was induced using the Nano-Glo Dual-Luciferase Reporter Assay System (Promega). Luciferase activity of firefly and Nano-Luc was evaluated as luminescence measured using a SpectraMax iD3 plate reader (Molecular Devices). Luciferase activity units were represented as the ratio of Nano-Luc to firefly luminescence.

### Quantitative real-time PCR

Total RNA was extracted using the RNAeasy Mini Kit (Qiagen), and cDNA was synthesized using SuperScript II reverse transcriptase (Invitrogen). The PCR reaction mixture was prepared on a 96-well plate using a Power SYBR Green PCR Master Mix according to the manufacturer’s instructions (Applied Biosystems). The primer sets are listed in Table S3. They were run in triplicate on a 7500 Fast Real-Time PCR System (Applied Biosystems). The expression level of each mRNA was estimated according to the corresponding standard curve and normalized to that of GAPDH. Data are displayed as percentages to a maximum value or relatives to an untreated sample.

**Table S3.**
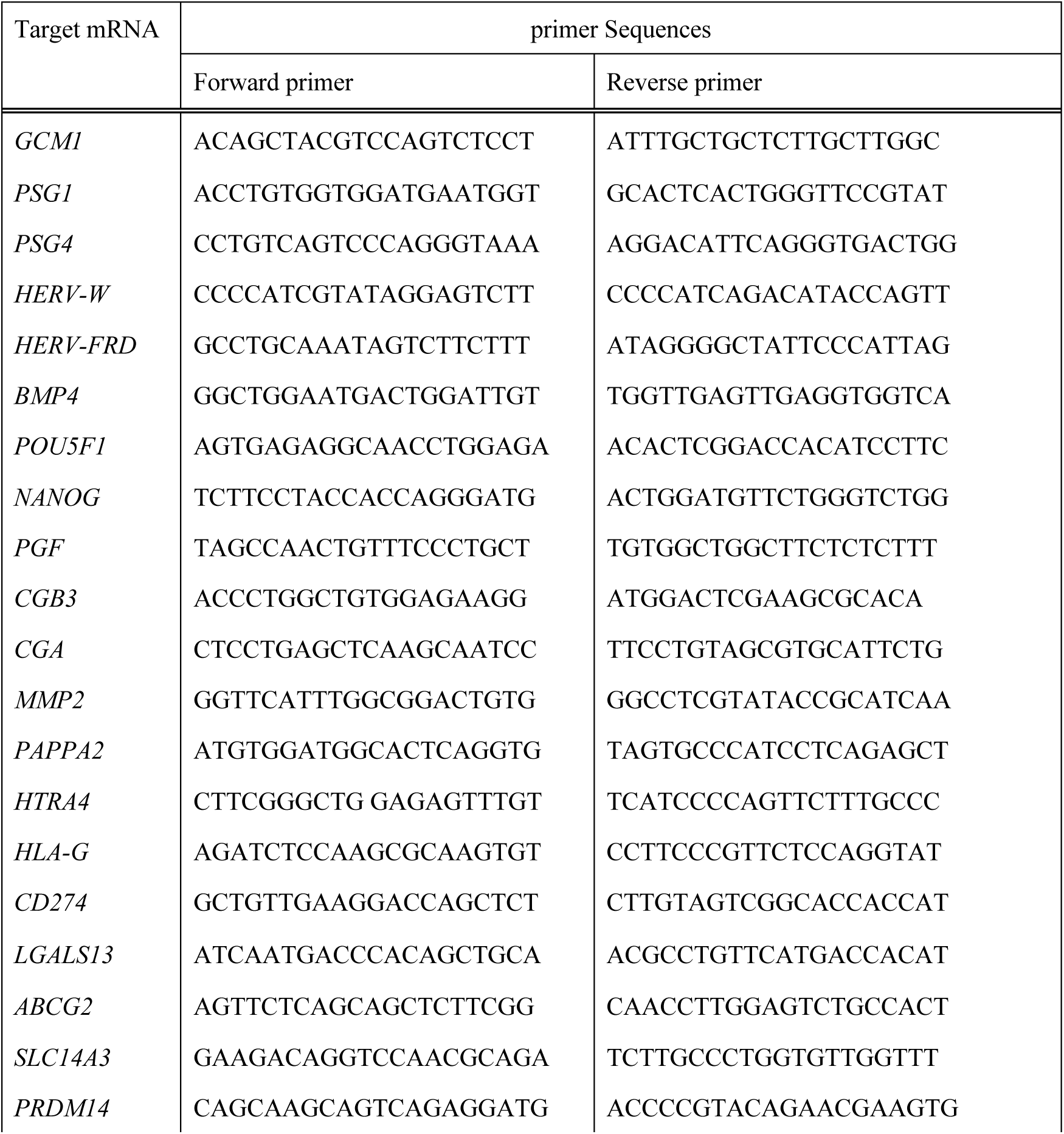
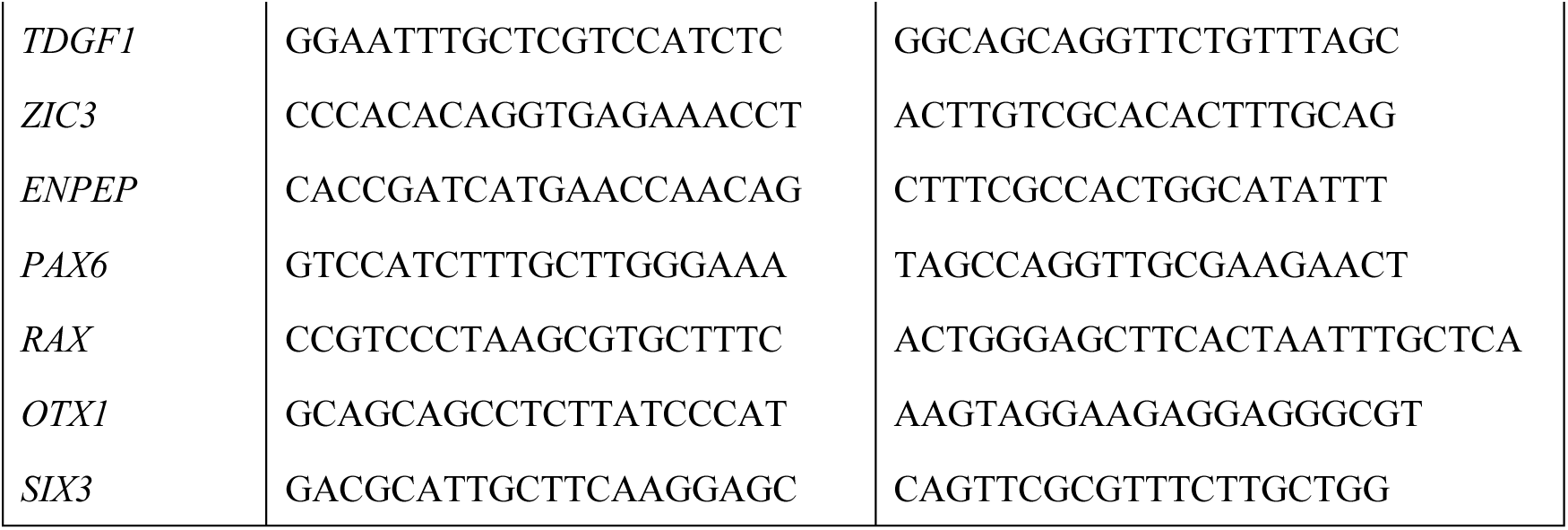
Primers used for quantitative RT-PCR

### Live imaging

Live imaging was performed using an inverted microscope (IX81-ZDC, Olympus) that was equipped with a stepper filter wheel (Ludl) and a cooled EM-CCD camera (ImagEM, Hamamatsu Photonics) as described previously (Ohgushi et al., 2015). For a confocal observation to monitor syncytium formation, a single slice of the image was recorded using a CSU-W1 unit (Yokogawa) configured with an IX81-ZDC microscope.

### Capture of syncytia formation

Two lines of KhES-1 cells expressing H2B-venus or mCherry were treated with AP for 4 days on the different culture plates. On day 4, these cultures were dissociated and mixed at 1 : 1 ratio. The mixed cells were seeded on Matrigel-coated 6-well plates (2.5 × 10^5^ cells per well) and further cultured in the DMEM/F12-based medium (D-MEM/F12 supplemented with 4% KSR additive, 1% ITS-X (Gibco), 0.3 % BSA (Invitrogen), and 0.1 mM 2-mercaptoethanol and 2.5 µM Y-27632). For live imaging, the mixed cells were seeded on Matrigel-coated 35 mm dishes (Idibi) and time-lapse recording was started on the next day. After 6 days of mixing, the cells were dissociated and harvested for flow cytometric analyses. To validate protein expression in syncytial cells, dissociated cells were suspended in a culture medium and stood in suspension for 1 h in a CO_2_ incubator. Double-positive cells (2 × 10^5^ cells) were collected using a FACS Aria IIIu. Western blot analyses were done as described above. For the comparison, undifferentiated ESCs (non-labeled) were subjected to a similar procedure.

### Bulk-RNA sequence and data analyses

Total RNA was harvested as described above. Sequencing libraries were prepared from 1 μg total RNA using the TruSeq Standard mRNA LT Sample Preparation Kit (Illumina) and sequenced by Illumina NextSeq500 (Illumina) using NextSeq500/550 High Output v2.5 Kit (Illumina) to obtain single-end 75 nt reads. Fastq files were generated using BaseSpace Onsite (Illumina). They are depositing to the Gene Expression Omnibus (GEO) database.

Sequence reads were aligned against GRCh38 genome assembly using STAR (version 2.6.1d). Read counts and transcripts per million (TPM) values of each gene were quantified and calculated using RSEM. The read counts and TPM values of each sample were imported into the R platform (version 4.0.3) as the matrix data. The genes for which the sum of read counts of all given samples was < 10 were excluded as ultra-low expressed genes for the following analyses. The read count matrix was provided to athe R package edgeR (version 3.32.0), and differential gene expression between the two samples was assessed using an exact test. The DEGs were defined as genes that were expressed with a *p*-value < 0.01 and a false discovery rate < 0.05. For gene ontology and tissue enrichment analyses, upregulated genes (log2[fold change] > 4) were extracted from DEGs and provided as input to the R package clusterProfiler (version 3.18.0) and TissueEnrich (version 1.10.1), respectively. TPM was used as a measure of relative transcript abundance. Hierarchical clustering was performed with the R function *dist* and *hclust* using ‘ward.D2’ based on TPM values. Principal component analyses were performed using the *prcomp* function in R based on the log-transformed TPM + 0.01 value.

### Single-cell RNA sequence and data analyses

The G3KI#2-5 hESCs were treated with AP for 4 days. They were dissociated by TrypLE select into single cells, and then the cell suspension was applied to a chromium controller (10x Genomics). A Chromium Single Cell 3’ Library & Gel Beads Kit v2 (10x Genomics) was used to generate oligo-dT-primed cDNA libraries following the manufacturer’s protocol. The cDNA library was then sequenced on an Illumina NextSeq500 (Illumina) using the NextSeq500/550 High Output v2.5 Kit (Illumina). The sequence reads were aligned against GRCh38 genome assembly using the *cellranger count* command of Cell Ranger (version 4.0.0) to generate a count matrix of unique molecular identifiers (UMIs) for each gene per cell. They are depositing to the GEO database. All subsequent analyses were done on the R platform using the Seurat package (version 3.2.3). The Seurat object was generated using genes expressed in at least five cells and cells including at least five genes. For quality control, cells that expressed < 1500 genes, > 4000 genes, and > 10 % of mitochondrial genes were filtered out. The final dataset contained 17,937 genes and 2,464 cells. The data were normalized and scaled using a *sctransform* function. The cell cycle effect was assessed using the *CellCycleScoring* function and was regressed by *vars.to.regress* option implemented in the *sctranform* function. All scale data for each cell are shown in Supplemental table 5. To reduce the dimensionality of the datasets, the *RunPCA* function was conducted using the top 3,000 highly variable genes. After nonlinear dimensional reduction was conducted with the *RunUMAP* function, cells were projected into a two-dimensional space using UMAP. We tested different numbers of PCs and selected the first 12 PCs for the *RunUMAP* function. Cell clusters were identified using the *FindClusters* function by the first 12 PCs with the resolution parameter set to 0.2. Signature scores for each cell were computed using the *AddModuleScore* function using the given genelist. To identify differentially expressed genes among the clusters of interest, we used the *FindMarkers* function. Statistical significance was tested using a Wilcoxon rank sum test, and genes with adjusted *p* values less than 0.01 were considered significant. For the trajectory inference and pseudotime analyses, we used the R packaged Monocle3 (version 0.2.3.0). The processed Seurat object was imported as a Monocle object using the *as.cell_data_set* function implemented in the SeuratWrappers package (version 0.3.0). We identified cell clusters and partitions using the *cluster_cells* function with the *k* value set to 8, and then fitted a principal graph using the *learn_graph* function. For pseudotime estimation, we defined the beginning root of the trajectory using the *order_cells* function considering the GATA3 expression level. Each cell was colored by the pseudotime and plotted on a UMAP space using the *plot_cells* function.

### External dataset

The expression data for single cells of human peri-implantation embryos were downloaded from Gene Expression Omnibus (GEO) with accession number GEO136447 as an FPKM matrix (Xiang et al., 2019). Gene lists for EPI, PrE, TrB, pre-CTB, post-CTB, early-STB, STB, early-EVT, and EVT were obtained from the report of Xiang et al. (Xiang et al., 2020, their Supplemental Table 1 and 4). Genes enriched in cynomolgus monkeys early and late AM were defined by Ma et al. The AM gene lists were obtained by extracting the genes annotated as ‘E-AM’ or ‘L-AM’ in the column of ‘cluster’ (Ma et al., 2019, their Table S6). These gene lists were converted to the human version using a one-to-one annotation table for human-cynomolgus monkey gene symbols (Nakamura et al., 2016, their Supplemental table 2). A gene set for the human placenta was generated by extracting the genes annotated as ‘Placenta’ in the column of ‘max_organ’ (Cao et al., 2020, their Table S2). AM-unique gene sets were generated by subtracting the placenta genes from the AM gene list. Other datasets used in this study were downloaded from GEO with accession numbers GSE101655 (Cliff et al., 2017), GSE150616 (Liu et al., 2020), GSE144994 (Io et al., 2021) and GSE138688 (Dong et al., 2019), or Japanese Genotype-phenotype Archive (JGA) with accession numbers JGAD00000000073 and JGAD00000000115 (Okae et al., 2019).

### Graphical presentation

Correlation heatmaps and gene expression heatmaps were generated using the R package corrplot (version 0.84) and heatmaply (version 1.1.1), respectively. Graphic visualization of single-cell transcriptome data was done using *DimPlot*, *FeaturePlot*, *VlnPlot*, and *DoHeatmap* functions of Seurat with some modifications using the R package ggplot2 (version 3.3.3). Flow cytometric data were visualized using the FlowJo software (version 10.7.1). All other graphs were generated using ggplot2.

### Statistical analysis

Error bars shown in the figures represent standard deviations and n in the legends is the number of experiments. Statistical significance (two-sided) was tested by Student’s t-test or Wilch t-test for two-group comparison, Tukey’s (among all groups), or Dunnett’s test (versus control) for multiple-group comparison.

## Notes

### Competing Interest Statement

The authors have declared no competing interest.

