## Supplemental Figures for "Cell-autonomous differentiation of human primed embryonic stem cells into trophoblastic syncytia through the nascent amnion-like cell state"

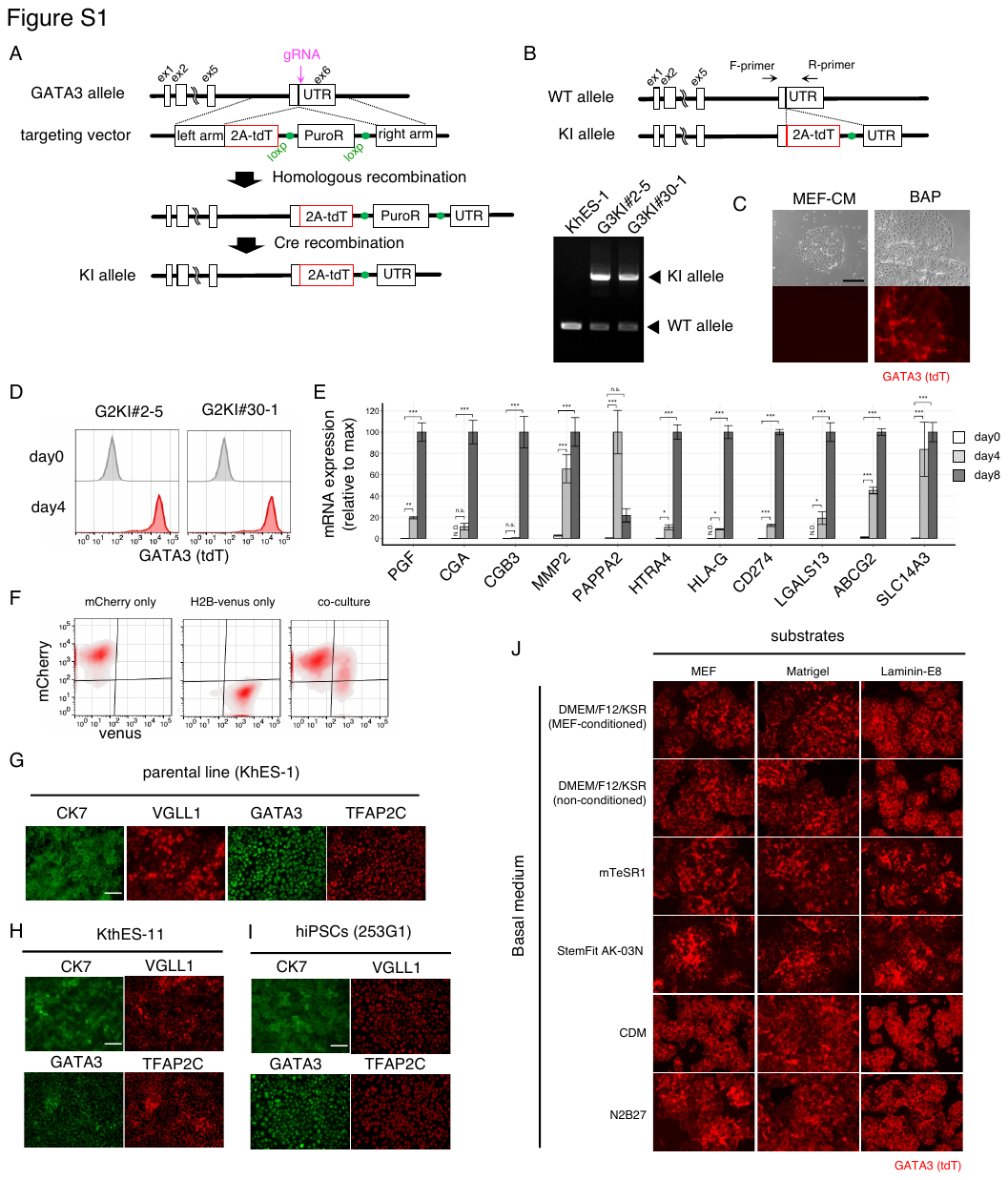

Figure S1. Trophoblast-like differentiation from human pluripotent stem cells.

(A) Schematic diagram for the generation of GATA3 reporter cell line.

(B) PCR genotyping of the GATA3-knockin allele. Corresponding regions for primers (F- or R-primer) were indicated. Genome extracted from parental KhES-1 and two independent knock-in lines were subjected to PCR analyses.

(C) Bright and fluorescent images of G3KI#2-5 cells treated with BAP for 4 days. Cells treated with MEF-CM for 4 days were shown as a control.

(D) Representative flowcytometric panels before and after treatment with AP. The results using two different clones were shown.

(E) qPCR assay for trophoblast-related genes. G3KI#2-5 cells were treated with AP for the indicated periods.

(F) Flowcytometric panels indicating expression level of H2B-Vunes or mCherry in differentiation cells at day10. Co-culture of Venus- or mCherry-expressing cells during differentiation produces dual positive cells (a red enclose in the right panel).

(G-I) Immunostaining for the indicated trophoblast markers in the parental cells (KhES-1, G), another ESC line (KthES-11, H) and an induced pluripotent cell line (253G1, I).

(J) Fluorescent images of G3KI#2-5 cells treated with AP for 4 days using different basal medium and substrates.

Scale bars show 100 µm. Error bars in the graphs represent standard deviations. Durnett’ test (n = 3) versus control (day 0) (); N. D., not detected, n. s., not significant; **p* <0.05, ** *p* <0.01, *** *p* < 0.001.

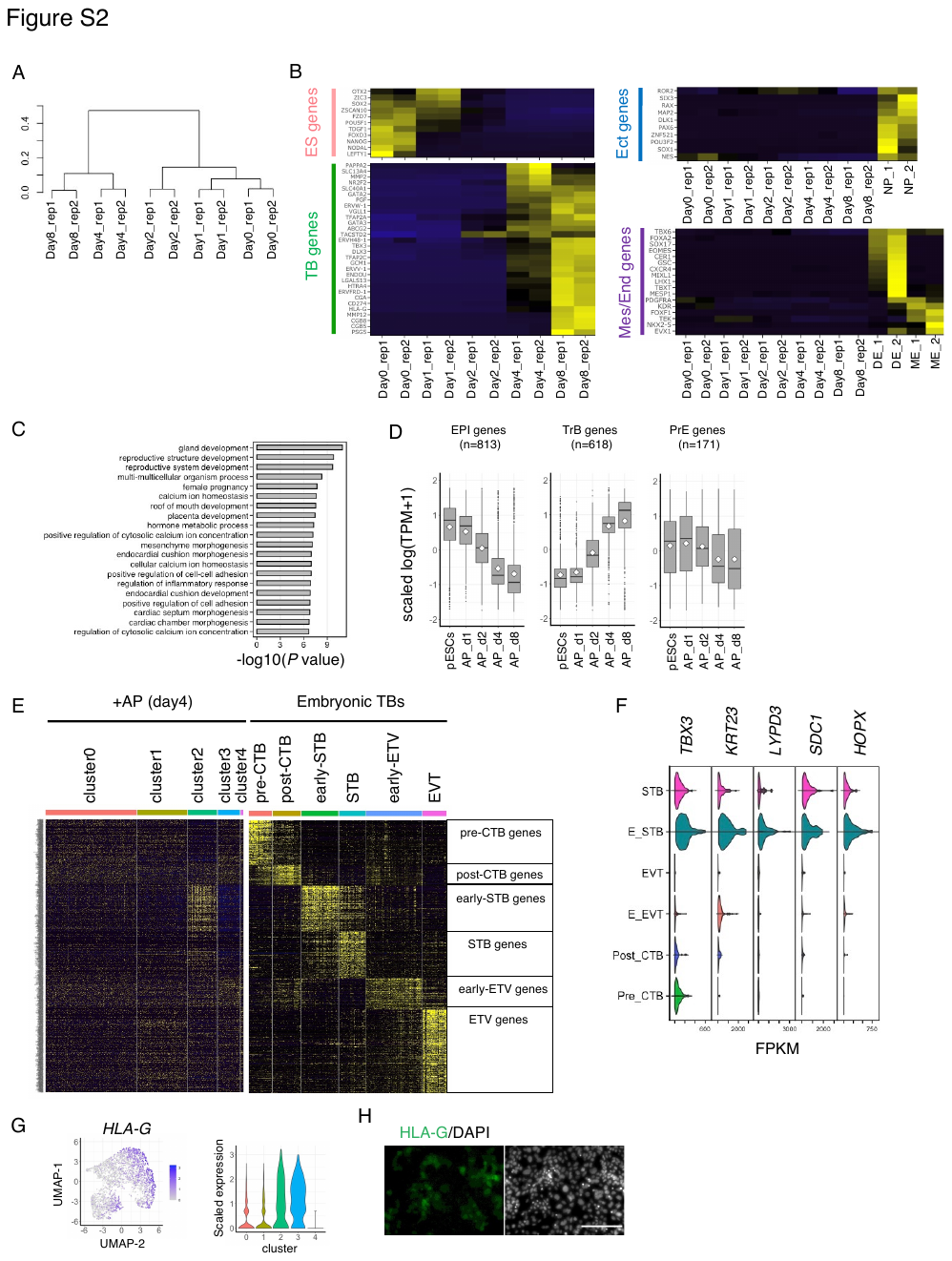

Figure S2. Transcriptome analyses of AP-induced differentiation.

(A) Unbiased hierarchical clustering of the transcriptome data. TheG3KI#2-5 cells were treated with AP for 0, 1, 2, 4 and 8 days with two biological replicates.

(B) Heatmap representation of the expression dynamics of typical markers for ESCs (ES genes), trophoblast (TB genes), ectoderm (Ect genes) and mesendoderm (Mes/End genes). The scaled TPMs value were used for gene expression. As positive control, TPMs from the published dataset were included (Neural progenitor; NP, definitive endoderm; DE, splanchnic mesoderm; ME, Cliff et al., 2017)

(C) Gene ontology analyses using the same geneset for Figure 2B.

(D) Boxplot presentation showing the expression dynamics of the marker genes for the embryonic epiblast (EPI genes; n = 813), trophoblast (TrB genes; n = 618) and primitive endoderm (PrE genes; n = 171) (Xiang et al., 2020).

(F) Heatmap representation of the indicated marker genes that were identified as the embryonic trophoblast sublineages (Xiang et al., 2020). The used genes were the same to the Figure 2G.

(G) Violin plot presentations of gene expression in human embryos (Xiang et al., 2020). The FPKM values of genes specific for the cluster 2 were shown.

(H) Violin and scatter plot for HLA-G.

(I) Immunostaining for HLA-G expression in G2KI#2-5 cells after 6 days of AP addition.

Error bars in the graphs represent standard deviations. Durnett’ test (n = 3) versus control (day 0) (); N. D., not detected, n. s., not significant; **p* <0.05, ** *p* <0.01, *** *p* < 0.001.

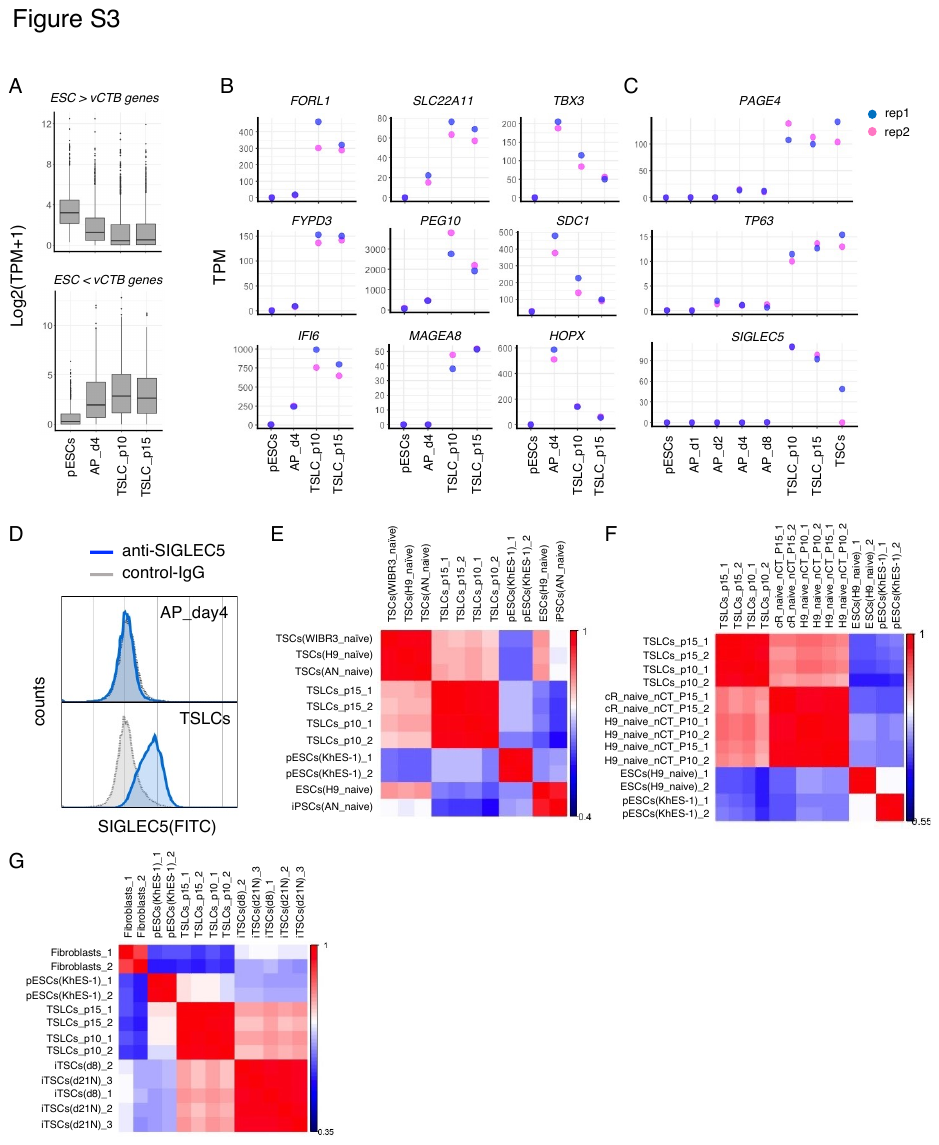

Figure S3. TSLC validation.

(A) TPM changes for the upregulated genes in ESCs relative to vCTB (ref) (FC >4, FDR > 0.01; n = 1,409) or in vCTB relative to ESCs (n = 857) were shown as boxplots (A).

(B) TPMs of the indicated genes in ESCs, AP-induced differentiated cells (day 4), TSLCs (passage number 10 and 15). Data were shown with two replicates.

(C) TPMs of the indicated genes in ESCs, AP-induced differentiated cells at four timepoints (day 1, 2, 4, 8), TSLCs (passage number 10 and 15). Data were shown with two replicates.

(D) Flowcytometric panel for TSLC-specific SIGLEC-5. Cells treated with AP for 4 days and TSLCs were stained with the indicated antibodies.

(E-G) Spearman correlation of transcriptomes from ESCs and TSLCs with published datasets. Comparison with dataset from Dong et al. (E), Io et al. (F) and Liu et al. (G) were shown.

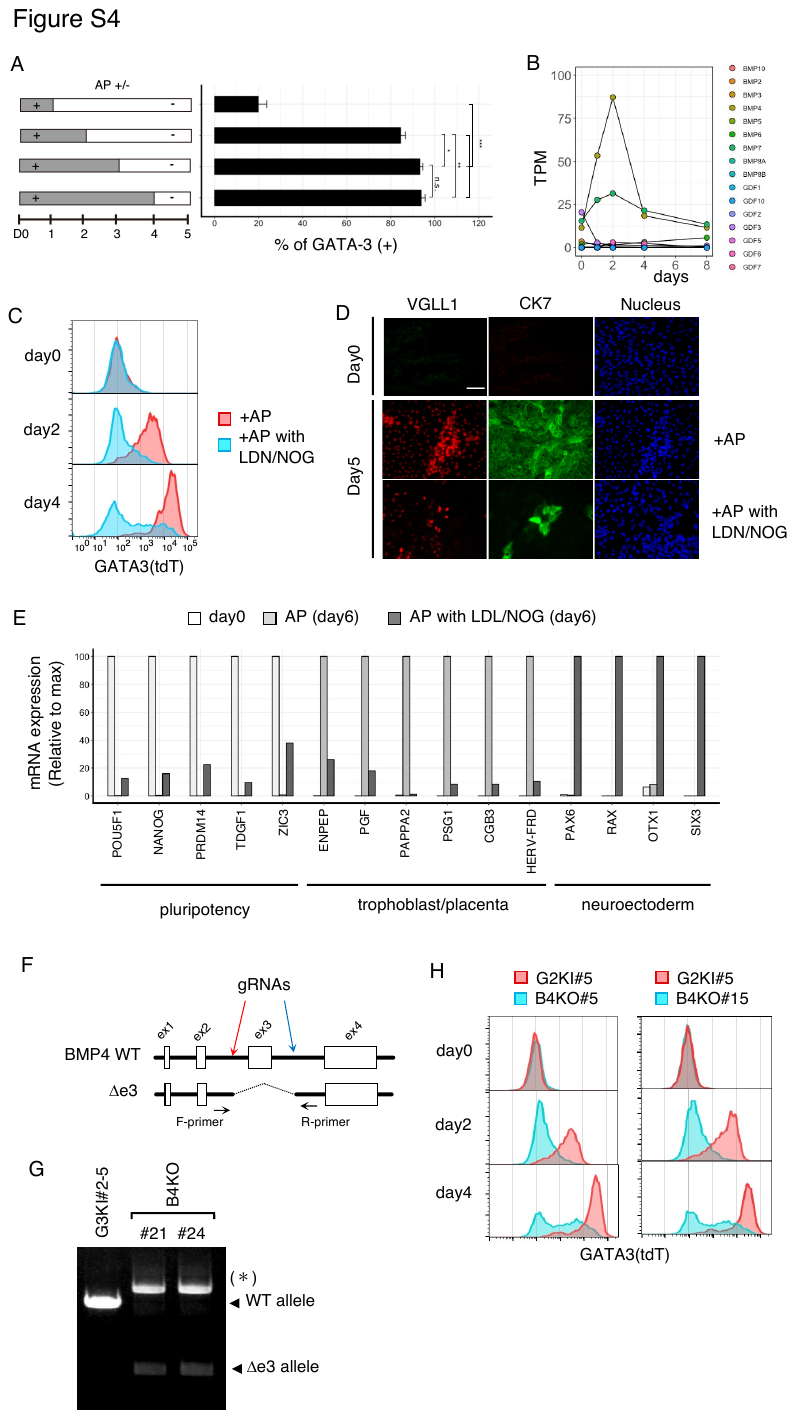

Figure S4. Mechanistic analyses of AP-induced differentiation.

(A) Time-window analysis of AP treatment on GATA3^+^ cell induction. AP was present during the periods indicated as gray bars. Flowcytometry analyses were performed at day 5.

(B) Expression dynamics for the indicated BMP ligands. Data was shown as average TPMs.

(C) Time course quantitation of GATA3^+^ cells. The G3KI reporter cells were treated with AP for the indicated periods in the presence or absence of BMP blockers.

(D) Immunostaining using KhES-1 cells for the indicated trophoblast markers. Scale bars show 100 µm

(E) qPCR assay for the indicated markers specific for pluripotent stem cells, trophoblasts and neuroectoderm. Cells were treated with AP in the presence or absence of BMP blockers for the BMP blockers.

(F-G) Target region of the designed gRNAs and corresponding regions for primers (F- or R-primer). Genome extracted from parental G3KI#2-5 and two independent knockout clones were subjected to PCR genotyping. Asterisk indicate a nonspecific band amplified by PCR reaction.

(H) Flow cytometric panels for GATA3-tdT expression of AP-treated BMP4 knockout cells.

Error bars in the graphs represent standard deviations. Tukey’ test (n = 3) for multiple comparisons (A). **p* <0.05, ***p* <0.01, ****p* < 0.001.

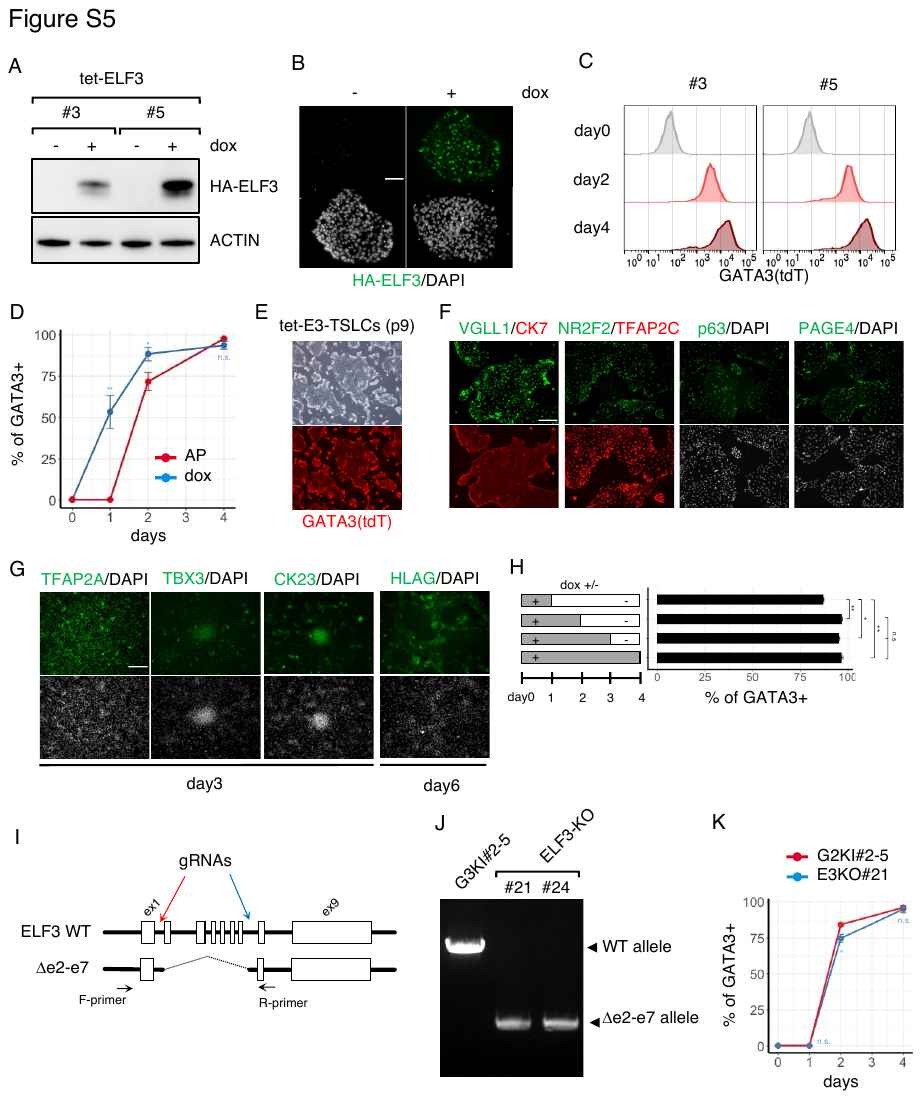

Figure S5. Characterization of trophoblast determinant ELF3.

(A) Expression of HA-ELF3 in dox-treated tet-ELF3 clones.

(B) Immunostaining analyses for expression and nuclear localization of HA-tagged ELF3.

(C) Flow cytometric panels for GATA3-tdT expression levels of dox-treated tet-E3 clones.

(D) Time course quantitation of GATA3^+^ cells. The tet-E3#3 clone was treated with AP (red) and dox (blue) for the indicated periods.

(E-F) TSLC derivation from dox-treated tet-E3#3 cells. Bright and red fluorescent images of live TSLCs were shown (E). They express the indicated TSC-related proteins (F). Scale bars show 100 µm.

(G) Immunostaining of dox-treated tet-E3#3 clone by the indicated trophoblast markers. Cells were treated with dox for 4 or 6 days. Scale bars show 100 µm.

(H) Time-window analysis of dox on the induction of GATA3^+^ cells. Dox was present during the periods indicated as gray bars. Flow cytometry analyses were performed at day4.

(I-J) Target region of the designed gRNAs and corresponding regions for primers (F- or R-primer) (I). Genome extracted from parental G3KI#2-5 and two independent knockout clones were subjected to PCR genotyping (J).

(K) Time course quantitation of GATA3^+^ cells. The E3KO clone and the parental cells were treated with AP for the indicated periods.

Error bars in the graphs represent standard deviations. Unpaired student’s t test (n = 3) at each time point (D, K); Tukey’ test (n = 3) for multiple comparisons (H). **p* < 0.05, ***p* < 0.01, ****p* < 0.001.

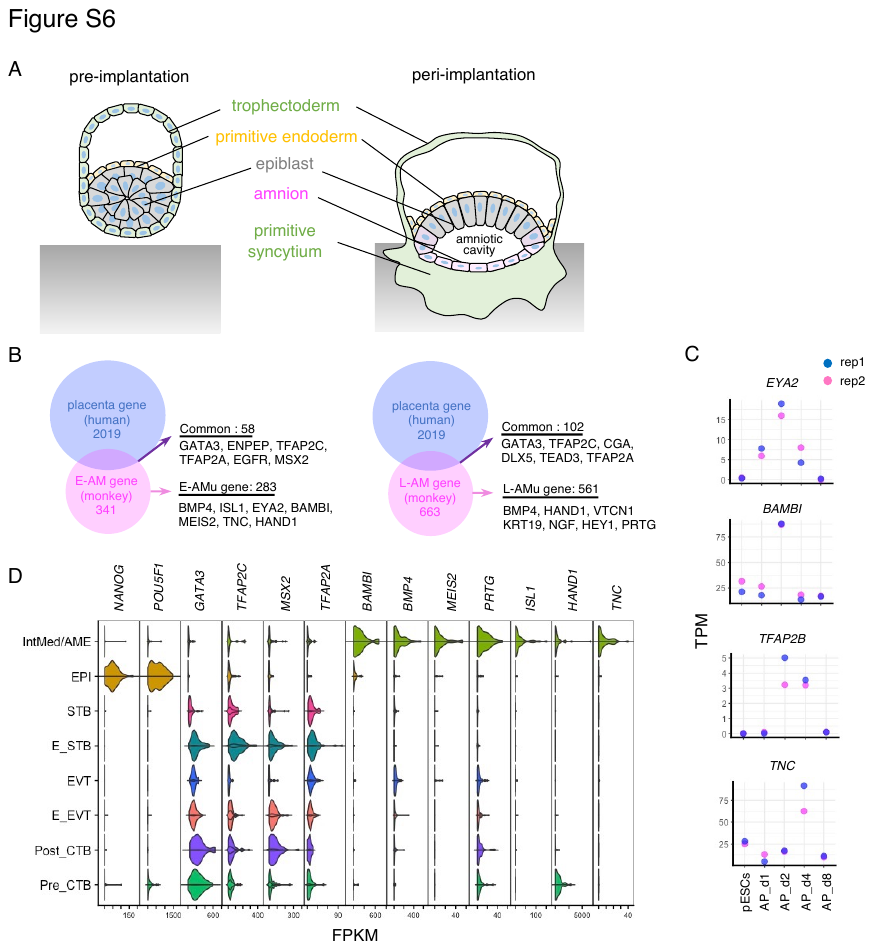

Figure S6. Nascent amnion-like features of AP-induced cells.

(A) Schematic diagram for pre- and post-implantation human embryo.

(B) Scheme for the selection of AM-unique geneset. The cynomolgous monkey AM gene list and the human placental gene list was obtained from the published literatures (Ma et al., 2019 for AM gene list: Cao et al., 2020 for placental gene list).

(C) TPMs of the indicated genes in ESCs and AP-induced differentiated cells. Data were shown with two replicates.

(D) Violin plot presentations of the indicated gene expression in EPI, trophoblast lineages (shown in Figure S2E) and amnion-like cells (IntMed/AME; intermediates and amnion) of human embryos (Xiang et al., 2020). The FPKM values of the indicated genes were shown.

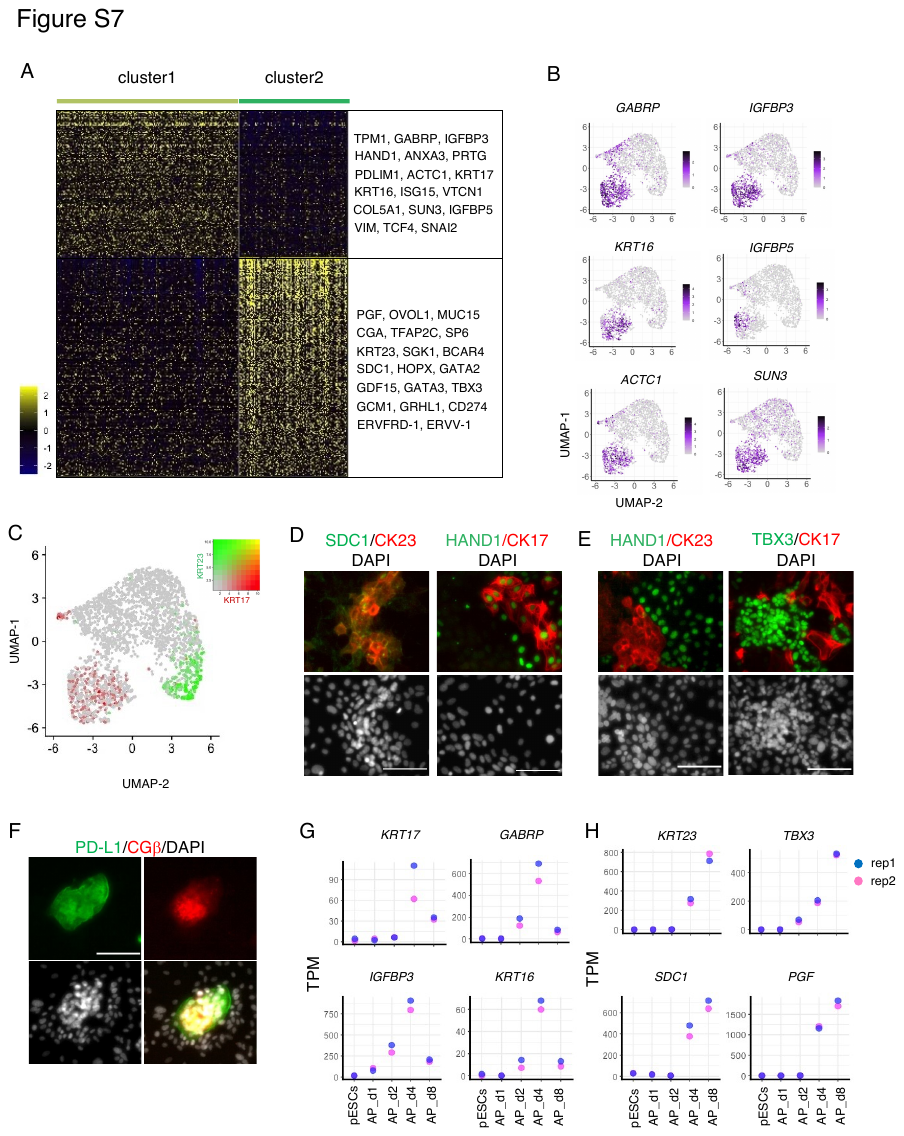

Figure S7. Co-existence of two extraembryonic cells within the same culture.

(A) Heatmap presentation for differentially expressing genes between the indicated clusters.

(B) Scatter plot representation of the cluster 1-enriched genes.

(C) Scatter plot for the colocalization scores of two cytokeratin. Red and green dots show expression level of cytokeratin 17 and 23 in each cell. Yellow dots, which show cells co-expressing these cytokeratins, were barely observed.

(D-E) Immunostaining of KhES-1 cells treated with AP for 5 days for the indicated AM- or STB-markers. Scale bars show 200 µm.

(F) Immunostaining of KhES-1 cells treated with AP for 8 days. Scale bars show 100 µm.

(G-H) TPM dynamics of the indicated AM genes (F) and STB genes (G) in ESCs and AP-induced differentiated cells. Data were shown with two replicates.
